# Resistor: an algorithm for predicting resistance mutations using Pareto optimization over multistate protein design and mutational signatures

**DOI:** 10.1101/2022.01.18.476733

**Authors:** Nathan Guerin, Andreas Feichtner, Eduard Stefan, Teresa Kaserer, Bruce R. Donald

## Abstract

Resistance to pharmacological treatments is a major public health challenge. Here we report Resistor—a novel structure- and sequence-based algorithm for drug design providing prospective prediction of resistance mutations. Resistor computes the Pareto frontier of four resistance-causing criteria: the change in binding affinity (Δ*K*_*a*_) of the (1) drug and (2) endogenous ligand upon a protein’s mutation; (3) the probability a mutation will occur based on empirically derived mutational signatures; and (4) the cardinality of mutations comprising a hotspot. To validate Resistor, we applied it to kinase inhibitors targeting EGFR and BRAF in lung adenocarcinoma and melanoma. Resistor correctly identified eight clinically significant EGFR resistance mutations, including the “gatekeeper” T790M mutation to erlotinib and gefitinib and five known resistance mutations to osimertinib. Furthermore, Resistor predictions are consistent with sensitivity data on BRAF inhibitors from both retrospective and prospective experiments using the KinCon biosensor technology. Resistor is available in the open-source protein design software OSPREY.

## 1. Introduction

Acquired resistance to therapeutics is a pressing public health challenge that affects maladies from bacterial and viral infections to cancer [1–6]. There are several different ways cancer cells acquire resistance to treatments, including drug inactivation, drug efflux, DNA damage repair, cell death inhibition, and escape mutations, among others [2]. Accurate, prospective prediction of resistance mutations could allow for design of drugs that are less susceptible to resistance. While it is unlikely that medicinal chemists will be able to address all of the resistance-conferring mechanisms in cancer cells, progress can be made by the incorporation of increasingly accurate models of the above contributing factors to acquired resistance, leading to the development of more durable therapeutics. To that end, several structure-based computational techniques for therapeutic design and resistance prediction have been proposed.

One such technique is based on the substrate-envelope hypothesis. In short, the substrate-envelope hypothesis states that drugs designed to have the same interactions as the endogenous substrate in the active site will be unlikely to lose efficacy because any mutation that ablates binding to the drug would also ablate binding to the endogenous substrate [7]. C. Schiffer and B. Tidor’s labs developed the substrate-envelope hypothesis for targeting drug-resistant HIV strains [7–10]. Their design technique has been successfully applied to develop compounds with reduced susceptibility to drug-resistant HIV proteases [10].

Another computational technique is to use ensemble-based positive and negative design [11, 12]. There are two specific ways that point mutations can confer resistance to therapeutics: they can decrease binding affinity to the therapeutic or they can increase binding to the endogenous ligand [11, 13]. Protein design with the goal of decreasing binding is known as *negative design*, and increasing binding is known as *positive design*. As a concrete example, consider the case of a drug that inhibits the tyrosine kinase activity of the epidermal growth factor receptor (EGFR) to treat lung adenocarcinoma. Here, an active site mutation could sterically prevent the inhibitor from entering the active site [14]. On the other hand, a different mutation might have no effect on an enzyme’s interactions with the drug but instead increase affinity to its native ligands, resulting in increased phosphorylization of downstream substrates [15, 16]. Because these two distinct pathways to therapeutic resistance exist, it is necessary to predict resistance mutations using both positive and negative design. In other words, predicting resistance can be reduced to predicting a ratio of the change in *K*_*a*_ upon mutation of the protein:endogenous ligand and protein:drug complexes.

*K*_*a*_ is an equilibrium constant measuring the binding and unbinding of a ligand to a receptor. It is defined as:

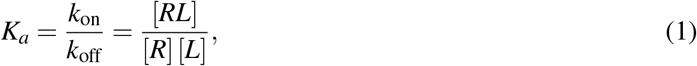

where *k*_on_ and *k*_off_ are the on- and off-rate constants, and [*RL*], [*R*], and [*L*] the equilibrium concentrations of, respectively, the receptor-ligand complex, unbound receptor, and unbound ligand. *K*_*a*_ is the reciprocal of the disassociation constant *K*_*d*_. *K*^*^ is an algorithm implemented in the OSPREY computational protein design software that provably approximates *K*_*a*_ [17, 18]. See the supplementary materials for further explanation of the *K*^*^ algorithm.

Our lab developed a provable, ensemble-based method using positive and negative *K*^*^ design to computationally predict and experimentally validate resistance mutations in protein targets [11]. We then applied this methodology to prospectively predict resistance mutations in dihydrofolate reductase when *Staphylococcus aureus* was treated with a novel antifolate [13] (also confirmed *in vivo* [13, 19]), demonstrating the utility of correctly predicting escape mutations during the drug discovery process. For cancer therapeutics, Kaserer and Blagg used OSPREY to combine positive and negative *K*^*^ design with mutational signatures and hotspot identification to both retrospectively and prospectively predict clinically significant resistance mutations [20]. Their technique combined sequence, in the form of trinucleotide mutational probabilities, and structures, in the form of positive and negative *K*^*^ design, to predict resistance mutations.

From these previous works, it is clear that multiple criteria must be combined to decide whether a mutation confers resistance. Often it is the human designers themselves who must choose arbitrary weights for different criteria. Yet multi-objective, or Pareto, optimization techniques would allow designers to combine multiple criteria without choosing arbitrary decision thresholds. Pareto optimization for protein design has been employed by Chris Bailey-Kellogg, Karl Griswold, and co-workers [21–26]. One such example is PEPFR (Protein Engineering Pareto FRontier), which enumerates the entire Pareto frontier for a set of different criteria such as stability vs. diversity, affinity vs. specificity, and activity vs. immunogenicity [27]. Algorithmically, PEPFR combined divide-and-conquer with dynamic or integer programming to achieve an algorithm where the number of divide-and-conquer “divide” steps required for the search over design space is linear only in the number of Pareto optimal designs. To our knowledge, Pareto optimization has yet to be applied to predicting resistance mutations, making Resistor the first algorithm to employ Pareto optimization for predicting resistance mutations.

Instead of merely finding a single solution optimizing a linear combination of functions, Pareto optimization finds all consistent solutions optimizing multiple objectives such that no solution can be improved for one objective without making another objective worse. Specifically, let Λ be the set of possible solutions to the multi-objective optimization problem, and let *λ* ∈ Λ. Let ℱ be a set of objective functions and *f* ∈ ℱ, where *f* : Λ→ ℝ is one objective function. A particular solution *λ* is said to *dominate* another solution *λ* ′ when

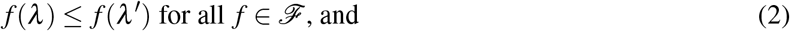

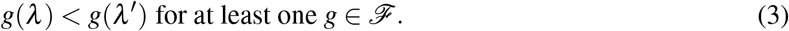

A solution *λ* is *Pareto optimal* if it is not dominated. Resistor combines ensemble-based positive and negative design, cancer-specific mutational signature probabilities, and hotspots to identify not only the Pareto frontier, but also the Pareto ranks of all candidate sequences.

The inclusion of mutational signature probabilities in Pareto optimization is possible because distinct mutational processes are operating in different types of cancers [28, 29]. Specifically, these mutational processes drive the type and frequency of DNA base substitutions. Each different signature is postulated to be associated with a biological process (such as ABOPEC activity [29]) or a causative agent (such as tobacco use), although not all associations are definitively known. What is certain is that particular signatures tend to appear in particular types of cancer. For example, 12 single-base substitution signatures, 2 double-base substitution signatures, and 7 indel signatures were found in a large set of melanoma samples, with many of those signatures associated with ultraviolet light exposure [29]. Building on the work of Alexandrov et al. [28], Kaserer and Blagg combined the multiple signatures found in each cancer type to generate overall single-base substitution probabilities [20]. Resistor uses these probabilities to compute the overall probability that mutation events will occur in a gene independent of changes to protein fitness. This amino acid mutational probability is one of the axes we optimize over.

The most computationally complex part of provable, ensemble-based multistate design entails computing the *K*^*^ scores of the different design states. This is largely because for biological accuracy it is necessary to use *K*^*^ with continuous sidechain flexibility [30, 31]. Though OSPREY has highly-optimized GPU routines for continuous flexibility [18], energy minimization over a combinatorial number of sequences in a continuous space is, in practice, computationally expensive. Having a method to reduce the number of sequences evaluated would greatly decrease the computational cost. Comets is an empirically sublinear algorithm that provably returns the optimum of an arbitrary combination of multiple sequence states [32]. Resistor uses Comets to prune sequences whose predicted binding with the drug improves and binding with the endogenous ligand deteriorates. While Comets does not compute the full partition function, it provides a useful method to efficiently prune a combinatorial sequence space. This is particularly important when one considers resistant protein targets with more than one resistance mutation. By virtue of pruning using Comets, Resistor inherits the empirical sublinearity characteristics of the Comets sequence search, rendering Resistor, to our knowledge, the first provable structure-based resistance prediction algorithm that is sublinear in the size of the sequence space.

The tyrosine kinase EGFR and serine/threonine-protein kinase BRAF are two oncogenes associated with, respectively, lung adenocarcinoma and melanoma. Both kinases are conformationally flexible, but two conformations are particularly determinative to their kinase activity—the “active” and “inactive” conformations. Oncogenic mutations to EGFR include L858R and deletions in exon 19, both of which constitutively activate EGFR [33, 34]. Likewise, V600E is the most prevalent constitutively activating mutation in BRAF [35]. Numerous drugs have been developed to treat the EGFR L858R and BRAF V600E mutations. The first generation inhibitors erlotinib and gefitinib competitively inhibit ATP binding in EGFR’s active site, whereas binding by the third generation osimertinib is irreversible [36–38]. For BRAF, the therapeutics dabrafenib, vemurafenib, and encorafenib were designed to target the V600E mutation and are in clinical use, and PLX8394 is in clinical trials [39–42]. Use of Resistor to predict resistance mutations to these drugs would provide strong validation of the efficacy of this novel approach.

By presenting Resistor, this article makes the following contributions:

1. A novel multi-objective optimization algorithm that combines four axes of resistance-causing criteria to rank candidate mutations.
2. The use of Comets as a provable and empirically sublinear pruning algorithm that removes a combinatorial number of candidate sequences before expensive ensemble evaluation.
3. A validation of Resistor that correctly predicted eight clinically significant resistance mutations in EGFR, providing explanatory ensemble-bound structural models for acquired resistance.
4. Prospective predictions with explanatory structural models and experimental validation of resistance mutations for four drugs targeting BRAF mutations in melanoma.
5. Newly modelled structures of EGFR and BRAF bound to their endogenous ligands and inhibitors in cases where no experimental structures exist.
6. An implementation of Resistor in our laboratory’s free and open source computational protein design software OSPREY [18].

## 2. Methods

The Pareto optimization in Resistor optimizes four axes: structure-based positive design, structure-based negative design, sequence-based mutational probabilities, and the count of resistance-causing mutations at a given amino acid location. Briefly, we chose these four criteria because they identify mutations that 1) increase affinity to the endogenous ligand in such a way that it outcompetes the inhibitor; 2) decrease the efficacy of the drug by reducing its binding (leading to the same effect); 3) are predicted to occur based on the DNA sequence and excludes those that are unlikely to arise; and, 4) are located at residue positions where many mutations are predicted to confer resistance, thus identifying a position of relative importance. We believe these criteria to be the minimal requirements a cancer clone must fulfill to confer resistance, and we’ve had success predicting retrospective and prospective resistance mutations in a previous study using these four criteria [20]. It should be mentioned that, as a generalizable method, additional resistance-causing criteria could be trivially added to Resistor for further refinement. The Pareto optimization objective function maximizes the Δ*K*_*a*_ of the positive design (the protein bound to the endogenous ligand), minimizes the Δ*K*_*a*_ of the negative designs (the protein bound to the drug), maximizes the mutational probability, and maximizes the count of resistance-causing mutations per amino acid. Positive design (affinity to the endogenous ligand) is also an indication of clonal fitness, i.e. whether the mutated protein can still provide the function that the cancer cell so critically depends on. If the binding to the endogenous ligand is disrupted, then it’s unlikely the clone will survive. Fig. 1 shows an overview how these axes are implemented in our algorithm.

**Fig. 1:**
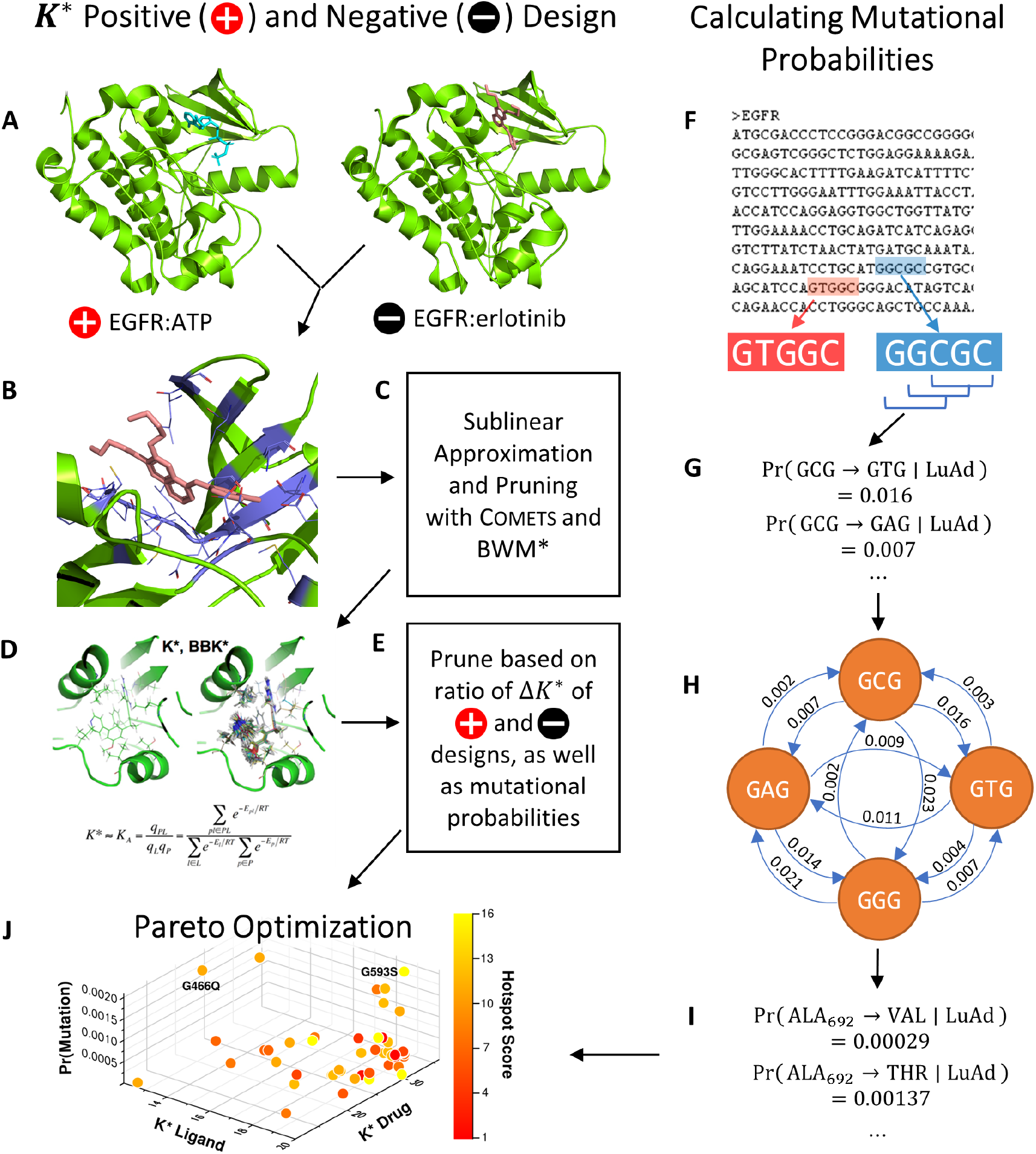
An example Resistor workflow with EGFR. Resistor finds the Pareto frontier from OSPREY positive and negative designs, mutational probabilities, and resistance hotspots. (A) Two structures are required as input to OSPREY to compute postive and negative design *K*^*^ scores. The structure for positive design is EGFR (green) bound to its endogenous ligand ATP (blue), for the negative design EGFR is bound to the drug erlotinib (pink). The goal of positive (resp. negative) design is to improve (resp. ablate) binding affinity. A mutation is resistant when its ratio of positive to negative *K*^*^ scores increases. (B) All residues within 5 Å (purple) of the drug are allowed to mutate to any other amino acid. (C) Comets is used as an efficient, sublinear algorithm to quickly prune infeasible mutations. BWM^*^ is used with a fixed branch width to compute a polynomial-time approximation to the *K*^*^ score. (D) Candidate mutations that pass the Comets pruning step have their positive and negative *K*^*^ scores computed in OSPREY. We recommend using the BBK^*^ with MARK^*^ algorithm as it is the fastest for computing *K*^*^ scores. (E) Candidate resistance mutations are pruned when their ratio of positive to negative *K*^*^ scores indicates a mutation does not cause resistance or if the target amino acid requires a mutation in all three DNA bases. (F) Resistor computes mutational probabilities using a protein’s coding DNA along with cancer-specific trinucleotide mutational probabilities for lung adenocarcinoma (abbreviated as LuAd), sliding a window (G) over 5′ - and 3′ -flanked codons. (H) Resistor employs a recursive graph algorithm to compute the probability that a particular amino acid will mutate to another amino acid (I). (J) Finally, Resistor uses Pareto optimization on the positive and negative *K*^*^ scores, the mutational probabilities, and hotspot counts to predict resistance mutants.

### 2.1. Structure-based Positive and Negative Design

We use the *K*^*^ algorithm in OSPREY to predict an *ε*-accurate approximation to the binding affinity (*K*_*a*_) in four states: 1) the wildtype structure bound to the endogenous ligand; 2) the wildtype structure bound to the therapeutic; 3) the mutated structure bound to the endogenous ligand; and 4) the mutated structure bound to the therapeutic. This *ε*-accurate approximation is called the *K*^*^ *score* [17, 18]. In order to calculate the *K*^*^ score of a protein:ligand complex, it is necessary to have a structural model of the atomic coordinates. Experimentally-determined complexes have been solved for EGFR bound to an analog of its endogenous ligand (PDB id 2itx), to erlotinib (1m17), gefitinib (4wkq), and to osimertinib (4zau) [43–46]. Similarly, we used the crystal structure for BRAF bound to dabrafenib (4xv2) and vemurafenib (3og7) [47, 48]. Experimentally-determined complexes of BRAF bound to encorafenib, PLX-8394, and an ATP analog in an active conformation do not exist, so we instead modelled the ligands into BRAF in its activated conformation (for additional details on model selection and preparation see the STAR methods). We used these predicted complex structures for our resistance predictions.

We added new functionality into OSPREY that simplifies the process of performing computational mutational scans. A *mutational scan* refers to the process of computing the *K*^*^ score of every possible amino acid mutation within a radius of a ligand. Resistor uses this functionality to calculate the four *K*^*^ scores for each amino acid within a 5 Å radius of the drug or the endogenous ligand. This generated a search space of 2471 sequences. We then set all residues with sidechains within 3 Å of the mutating residue to be continuously flexible for the Resistor *K*^*^ designs. Each sequence has an associated conformation space size dependent on the total number of mutable and flexible residues, which one can use as a heuristic to estimate the difficulty of computing a complex’s partition function. The average conformation space size of each sequence was ∼ 5.9 × 10^10^ conformations, thus computing the partition functions is only possible using OSPREY’s pruning and provable *ε*-approximation algorithms [18, 30, 49]. Empirical runtimes of the positive- and negative-*K*^*^ designs are shown in supplementary figure S1. The change in the *K*^*^ score upon mutation for the endogenous ligand (positive design) and drug (negative design) become two of the four axes of optimization. These two axes also form the basis of a pruning step (described in Sec. 2.4).

### 2.2. Computing the Probability of Amino Acid Mutations

To convert the trinucleotide to trinucleotide probabilities into amino acid to amino acid mutational probabilities, Resistor constructs a directed graph with the trinucleotides as nodes and the probability that one trinucleotide mutates into another trinucleotide as directed edges. It then reads the cDNA of the protein in a sliding window of 5′ - and 3′ -flanked codons, since the two DNA bases flanking a codon are necessary to determine the probabilities of either the first or third base of a codon mutating. We designed a recursive algorithm to traverse the graph and find all codons that can be reached within *n* single-base mutations, where *n* is an input parameter. The algorithm then translates the target codons into amino acids and, as a final step, sums the different probabilities on each path to an amino acid into a single amino acid mutational probability (see Fig. 1F-I). One can either (a) precompute a cancer-specific codon-to-codon lookup table consisting of every 5′ - and 3′ -flanked codon to its corresponding amino acid mutational probabilities, or (b) read in a sequence’s cDNA and compute the mutational probabilities on the fly. The benefit of (a) is it only needs to be done once per cancer type and can be used on an arbitrary number of sequences. On the other hand, when assigning mutational probabilities to proteins that have strictly fewer than 4^5^ amino acids, it is faster to compute the amino-acid specific mutational signature on the fly. In both cases, the algorithm is strictly polynomial and bounded by *O*(*kn*^9^), where *k* is the number of codons with flanking base pairs (upper-bounded by 4^5^) and *n* is the number of mutational steps allowed, which in the case of Resistor is 2. An implementation of this algorithm is included in the free and open source OSPREY GitHub repository [18].

### 2.3. Identifying Mutational Hotspots

After calculating the positive and negative change in affinity Δ*K*_*a*_ and determining the mutational probability of each amino acid, Resistor prunes the set of candidate mutations (see section 2.4). Post-pruning, it counts the number of mutations at each amino acid location. This count is important to determine whether a residue location is likely to become a “mutational hotspot”, namely a residue location where many mutations are predicted to confer resistance. Correctly identifying mutational hotspots is vital because they indicate that a drug is dependent on the wildtype identity of the amino acid at that location, and it is likely that many mutations away from that amino acid will cause resistance. Consequently, the fourth axis used in Resistor’s Pareto optimization is the count of predicted resistance-conferring mutations per residue location.

### 2.4. Reducing the Positive Prediction Space

Prior to carrying out the multi-objective optimization to identify predicted resistance mutations, we prune the set of candidates. First, we introduce a cut-off based on the ratio of *K*^*^ scores of positive and negative designs, adapted from Kaserer and Blagg’s 2018 cut-off [20]. We determine the average of the *K*^*^ scores for the drug and endogenous ligand across all of the wildtype designs for the same protein. The cut-off *c* is:

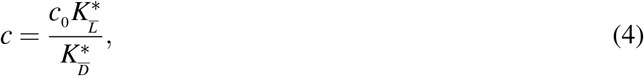

where *c*_0_ is a user-specified constant, 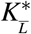 is the average of the *K*^*^ scores for the wildtype protein bound to the endogenous ligand, and 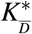 is the average of the *K*^*^ score for the wildtype protein bound to the drug. We recommend in practice to set *c*_0_ to be greater than the range 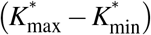 of wildtype *K*^*^ scores—we set it to for the tyrosine kinase inhibitor (TKI) predictions.^1^ A mutation *m* is predicted to be *resistant* when:

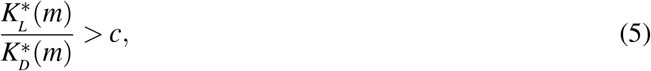

where 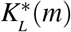 is the *K*^*^ score of the endogenous ligand bound to the mutant, and 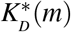 is the *K*^*^ score of the drug bound to the mutant.

We further prune the predicted resistance mutation candidates by removing all mutations that cannot arise within two DNA base substitutions. Whether an amino acid can be reached within two DNA base substitutions is determined by the algorithm described in section 2.2, and if it cannot, then that particular mutation is assigned a mutational probability of 0 and pruned.

## 3. Results

### 3.1. Resistor *Identifies 8 Known Resistance Mutations in EGFR*

We evaluated a total of 1257 sequences across the three TKIs for EGFR. Among these sequences, the average conformation space size for computing a complex’s partition function was ∼ 1.3 × 10^7^. After we ran the Resistor algorithm on these sequences, a total of 108 mutants were predicted as resistance-conferring candidates for all three inhibitors combined from a purely thermodynamic and probabilistic basis, i.e. these mutations were required to lower affinity of the drug in relation to the endogenous ligand (*K*^*^ Positive and Negative Design, Fig. 1A-D) and could be formed in patients by less than three base pair exchanges (Calculating Mutational Probabilities, Fig. 1F-I). To further prioritize mutations and identify those that are most likely to be clinically relevant, we then computed the Pareto frontier over the four axes for each drug (Fig. 1J). Remarkably, out of these 108 candidates, Resistor correctly prioritized eight clinically significant resistance mutants, with 7 of the 8 in the Pareto frontier of the corresponding inhibitor and the remaining mutant in the 2^nd^ Pareto rank (see Table 1). A detailed description of the result for each inhibitor is included in the sections below.

**Table 1:**
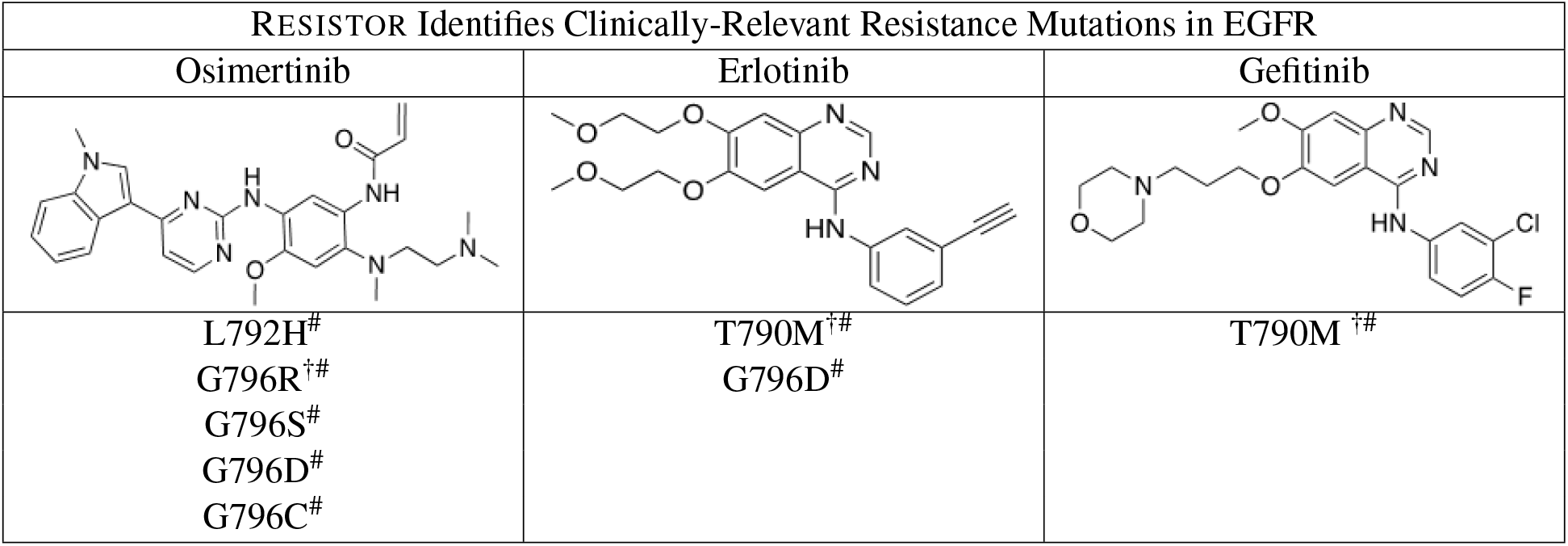
Resistor correctly identified 8 resistance mutations in EGFR to erlotinib, gefitinib, and osimertinib. For osimertinib, G796R, G796S, G796D, and G796C were on the Resistor-identified Pareto frontier. L792H was in the 2^nd^ Pareto rank. For erlotinib, both T790M and G796D were on the Pareto frontier. For gefitinib, T790M was also on the Pareto frontier. Previous studies have documented all of these resistance mutations as occurring in the clinic [50–57].† indicates that Resistor predicted the mechanism of resistance to be improved binding of the endogenous ligand to the mutant. # indicates that Resistor predicted the mechanism of resistance to be decreased binding of the drug to the mutant. Note that these predicted mechanisms are only attributed here if the predicted change in the log_10_(Δ*K*^*^) ≥ 0.5.

| RESISTOR Identifies Clinically-Relevant Resistance Mutations in EGFR |  |  |
| --- | --- | --- |
| Osimertinib | Erlotinib | Gefitinib |
| L792H<sup>#</sup><br>G796R<sup>†#</sup><br>G796S<sup>#</sup><br>G796D<sup>#</sup><br>G796C<sup>#</sup> | T790M<sup>†#</sup><br>G796D<sup>#</sup> | T790M<sup>†#</sup> |

#### 3.1.1. EGFR Treated with Erlotinib and Gefitinib

Of the 462 sequences evaluated for the TKI erlotinib, Resistor identified 50 as candidate resistance mutations. Pareto ranking placed 19 sequences on the frontier, 13 sequences in the second rank, and 11, 6, and 1 sequences in the third, fourth, and fifth ranks, respectively. Resistor correctly identified two clinically significant mutations, T790M and G796D, as being on the Pareto frontier [50, 51]. This is encouraging because T790M is, by far, the most prevalent resistance mutation that occurs in lung adenocarcinoma treated with erlotinib [58]. Similarly, for gefitinib, Resistor evaluated 438 sequences and identified 22 as candidate resistance mutants. The most relevant clinical mutant, T790M, is found on the Pareto frontier.

#### 3.1.2. EGFR and Osimertinib

Resistor evaluated 357 OSPREY-predicted structures of EGFR bound with osimertinib and EGFR bound with its endogenous ligand. Of those, 36 were predicted as resistance candidates. Pareto optimization placed 16 sequences on the frontier, 2 sequences in rank 2, 8 sequences in rank 3, 1 sequence in rank 4, and 5 sequences in rank 5. Resistor correctly identified five clinically significant resistance mutations to osimertinib: L792H, G796R, G796S, G796D, and G796C [52–57], and while L792H was in the 2^nd^ Pareto rank, all of the other correctly predicted resistance mutations are on the Pareto frontier.

Two osimertinib resistance mutations in particular stand out: L792H and G796D (see Fig. 2). Both of these mutants have appeared in the clinic [52–54, 57]. OSPREY generated an ensemble of the bound positive and negative complexes upon mutation, providing an explanatory model for how resistance occurs. In both cases, the mutant sidechains are much bulkier than the wildtype sidechain (Fig. 2A and D) and thus are predicted to clash with the original osimertinib binding pose (Fig. 2B and E). Consequently, in both cases the ligand is predicted to translate and rotate to create additional space for the mutant sidechains (Fig. 2C and F). We hypothesize that this movement weakens the other molecular interactions osimertinib makes in the EGFR active site.

**Fig. 2:**
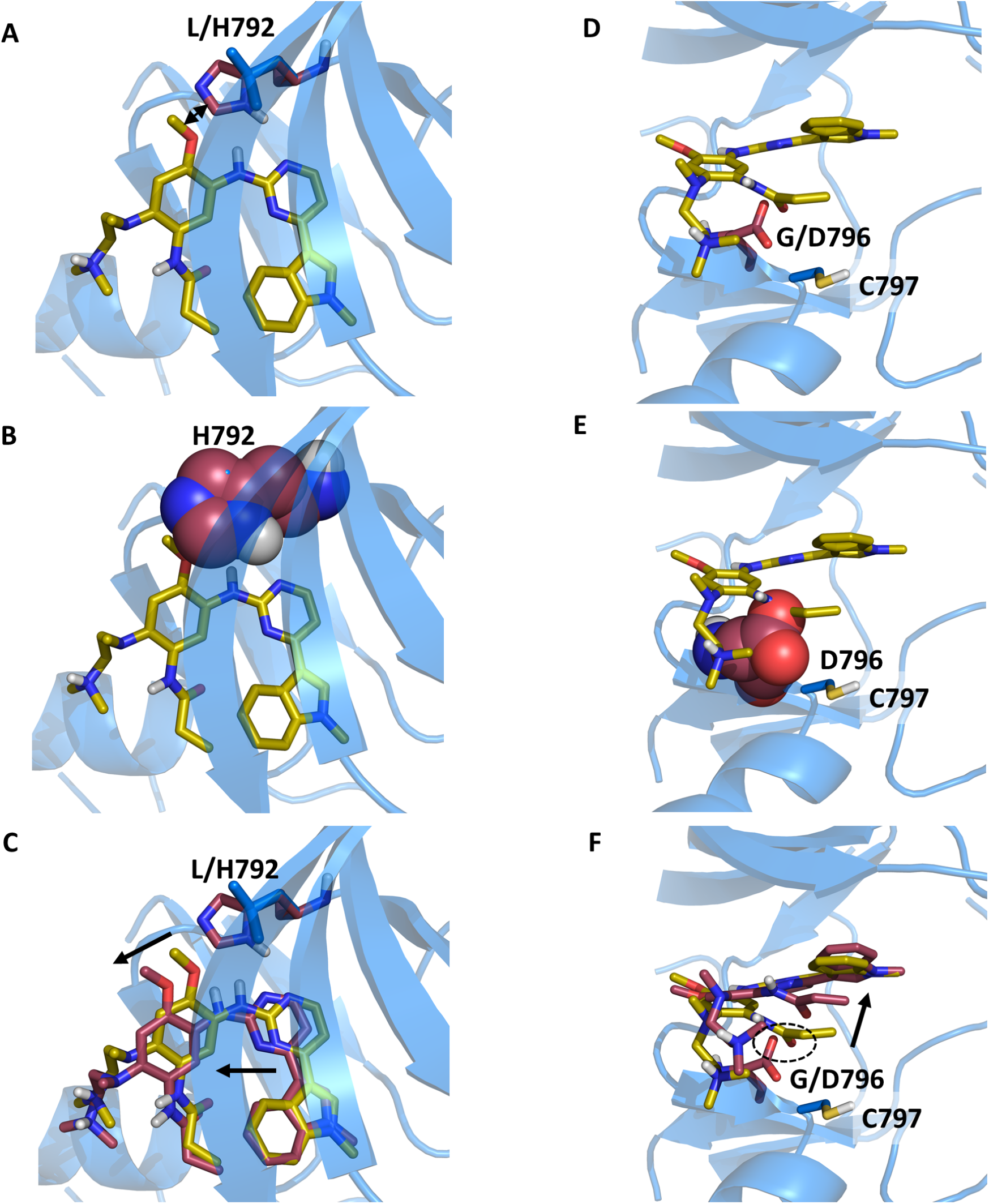
Structural models predicted by OSPREY agree with experimental data and explain mechanisms of Osimertinib resistance to EGFR mutations L792H and G796D. Structural models predicted by OSPREY of EGFR wildtype (blue) and resistance mutations (red) bound to osimertinib (yellow sticks). The histidine (A) and glutamate (D) side chains (red sticks) in the EGFR L792H (A) and G796D (D) mutations are bulkier than the wildtype leucine (A) and glycine (C) residues (blue sticks). They clash with osimertinib in its original binding pose as highlighted by the sphere representation in panels B and E. (C+F) To allow for accommodation of osimertinib in the modelled EGFR mutant structures (red sticks), the inhibitor’s position within the binding pocket moves from the experimentally determined binding pose (yellow sticks). Movements are indicated by black arrows. (F) In case of the G796D mutation, the carboxylate moiety of D796 is predicted to be in close proximity to the osimertinib amide oxygen (highlighted with the dashed circle), thus leading to electrostatic repulsion. This mutation site is adjacent to C797, which reacts with the allyl-moiety of osimertinib to form a covalent bond in the wildtype. Due to the steric and electrostatic properties of the G796D mutant, the allyl group is located further away from C797 in the model, thus preventing covalent bond formation. The movement of the allyl group is indicated by the black arrow.

In the case of G796D, there are additional factors that contribute to acquired resistance. First, the mutation to aspartate introduces a negative charge, which probably leads to electrostatic repulsion with the carbonyl oxygen of the osimertinib amide (Fig. 2F, highlighted with a dashed oval). In addition, the exit vector of the hydrogen bound to the amide nitrogen does not allow a hydrogen bond with the aspartate. Second, the allyl-group of osimertinib must be in close proximity to C797 for covalent bond formation. In fact, C797 is so important to osimertinib’s efficacy that mutations at residue 797 confer resistance [59, 60]. Even if osimertinib still binds to G796D, the allyl group would have to move away from C797 (Fig. 2F, highlighted with a black arrow). This would prevent covalent bond formation and thus reduce the efficacy of osimertinib considerably. Lastly, it is likely that the mutation away from glycine reduces the conformational flexibility of the loop, incurring an entropic penalty while also plausibly making it more difficult to properly align osimertinib and C797.

### 3.2. Resistor *Predicts New Resistance Mutations in BRAF and Provides Structural Models*

In addition to retrospective validation by comparison to existing clinical data for EGFR, we used Resistor to predict how mutations in the BRAF active site could confer resistance. Specifically, we used Resistor to predict which of 1214 BRAF sequences would be resistant to four TKIs—vemurafenib, dabrafenib, encorafenib, and PLX8394. On the Pareto frontier for vemurafenib are 13 mutations, for dabrafenib 16 mutations, for encorafenib 15 mutations, and for PLX8394 15 mutations. The full sets of predictions are included in the supplementary tables S4-S7. To validate Resistor’s predictions, we compared its predictions with two sources of experimental data: a saturation mutagenesis variant effect assay from Wagenaar et al. [61] and a cell-based kinase conformation reporter assay termed KinCon by Stefan and colleagues [62, 63]. Furthermore we carried out new KinCon experiments on a number of Resistor predictions to validate Resistor’s predictive capabilities.

#### 3.2.1. Retrospective and prospective validation of Resistor predictions using the BRAF KinCon biosensor reporter

KinCon, developed by Stefan and colleagues, is an in-cell protein-fragment complementation assay (PCA) that provides a readout of the activity conformation change of full-length BRAF upon mutation or exposure to different inhibitors [64]. KinCon’s bioluminescence assay functions by appending parts of a luceriferase enzyme to the N- and C-termini of full-length BRAF and observing the amount of bioluminescence, indicating whether BRAF favors an open, catalytically active or a closed, autoinhibited conformation (see Fig. 3A) [64]. Stefan and colleagues have demonstrated that activation of BRAF either via upstream regulators such as EGFR and GTP activated Ras or via tumorigenic mutations cause BRAF to favor an open conformation [62, 63]. The inhibitors bind to BRAF in the ATP binding site and cause BRAF’s N- and C-termini to interact, shifting BRAF back towards a more closed, intermediate state (see Fig. 3A) [62–64]. This implies that for inhibitor binding and BRAF closing to occur, a mutation (or a combination of mutations and/or upstream signaling events) needs first to induce an open conformation. Not all clinically observed BRAF mutations cause opening, even if they activate the MAPK pathway (e.g. L472C) [63, 65]. In the same vein, not all BRAF resistance mutants show increased kinase activity, in fact several are classified as kinase impaired [63, 65, 66]. One prominent mutation that shows both increased kinase activity and induces an open conformation is V600E (Fig. 3B). Inhibitor treatment shifts the V600E conformational equilibrium towards a more closed state [62, 63]. In contrast, the gatekeeper mutations T529M and T529I do not confer opening of the kinase conformation and are thus insensitive to inhibitor treatment [62]. However, in combination with V600E these mutations do confer resistance to BRAF inhibitors to varying degrees. Given that we model a state that is permissive of ligand binding at the outset (i.e., the ligand-bound BRAF complex), our Resistor calculations align very well with the reported KinCon measurements of double mutants (e.g. V600E/T529M and V600E/T529I).

**Fig. 3:**
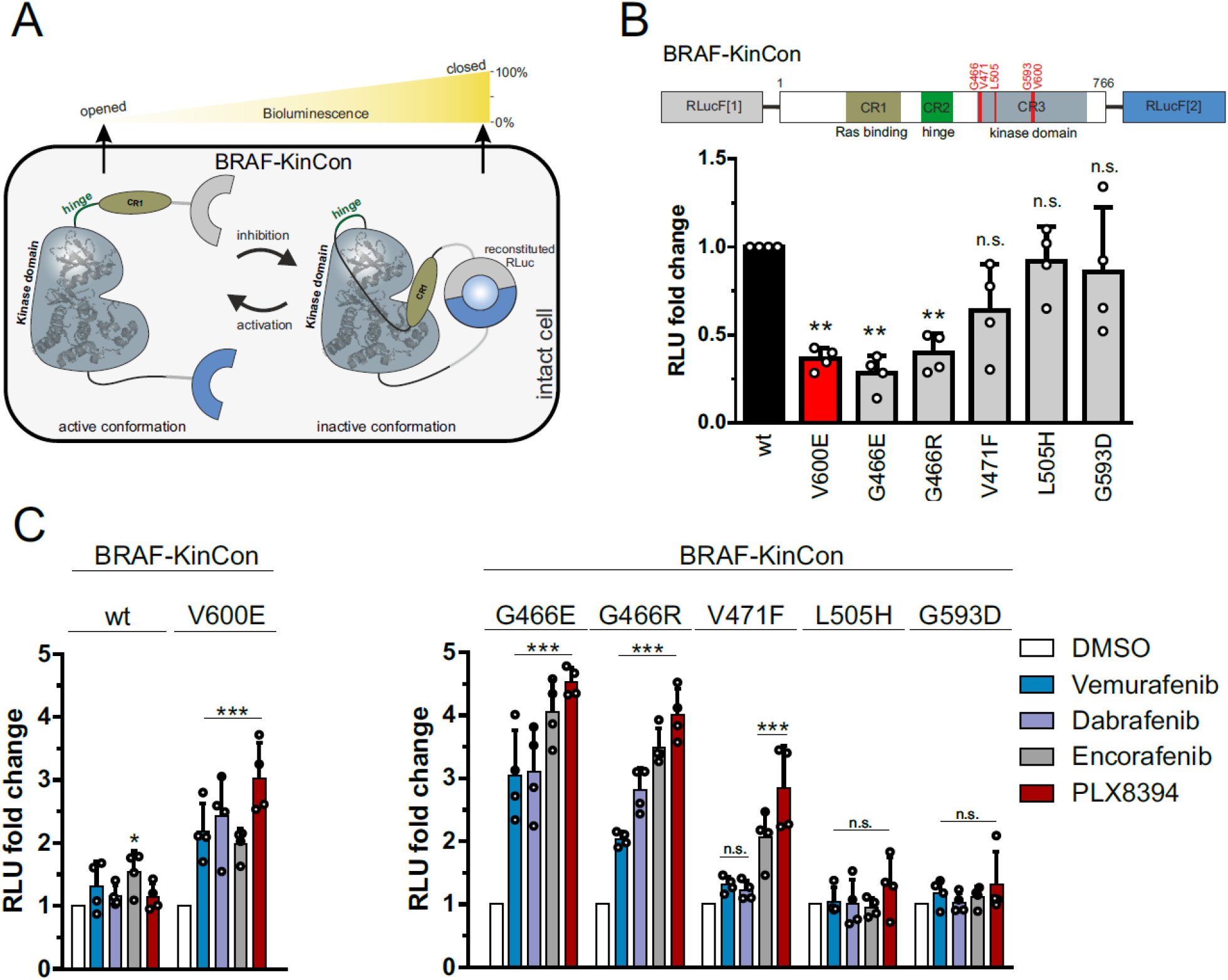
(A) Schematic depiction of Renilla luciferase (RLuc; F1: fragment 1, F2: fragment 2) PCA-based BRAF kinase conformation (KinCon) reporter system. Conformational rearrangement of the reporter upon (de)activation of the kinase are indicated. Closed kinase conformation induces complementation of RLuc PCA fragments resulting in increased RLuc-emitted bioluminescence signal. (B) Domain organization of the BRAF-KinCon reporter (top) and basal bioluminescent signals of the BRAF-wt (black), V600E (red), and novel mutant (grey) KinCon biosensors. Bars represent the mean signals, relative to BRAF-wt, in relative light units (RLU) with SD of four independent experiments (nodes). Raw bioluminescence signals were normalized on reporter expression levels, determined through western blotting. Asterisk indicates level of significance versus the wild type BRAF biosensor. (C) BRAF-KinCon biosensor dynamics, induced via treatment with respective BRAFi (1μM for 1h) prior to bioluminescence measurement. BRAF wt and V600E KinCon variants serve as control (left). The novel mutants are shown in a separate bar chart (right). Bars represent the mean signals, relative to the DMSO control, in relative light units (RLU) with SEM of four independent experiments (nodes). All experiments were performed in HEK293T cells 48 hours post transfection. \**p <* 0.05; \*\**p <* 0.01; \*\*\**p <* 0.001; n.s., not significant by t-test.

Specifically, the Resistor predictions of resistance concord with the previous KinCon biosensor results for V600E/T529M and V600E/T529I for three of the four inhibitors: vemurafenib, dabrafenib, and PLX8394 [62]. In the case of vemurafenib treatment, the proportion of open to closed conformations in the V600E/T529I mutant is not significantly different from the untreated V600E mutant, indicating vemurafenib treatment is not closing the conformational distribution in the double mutant [62]. These data agree with the Resistor calculation of the ratios of the log_10_ *K*^*^ scores, which predict that both double mutants are resistant to vemurafenib, with V600E/T529M more resistant. Treatment of BRAF with PLX8394 follows the same pattern as vemurafenib, namely the V600E/T529I mutant’s closed population increases only 1.2 fold compared to the untreated mutant, and the PLX8394-treated V600E/T529M mutant does not noticeably alter the conformational distribution [62]. In contrast, the PLX8394-treated V600E mutant’s closed population increases ∼3 fold compared to the untreated population, indicating V600E sensitivity to PLX8394 (see Fig. 3C). Resistor correctly predicted the V600E/T529I and V600E/T529M double mutants are resistant to PLX8394, with the change in the ratio of the log_10_ *K*^*^ scores of the two mutants suggesting that V600E/T529M confers greater resistance. In the case of dabrafenib, treatment of the V600E/T529I mutant closed the conformational distribution (2.4 fold more closed compared to untreated) more than treatment of the V600E mutation (2 fold more closed compared to untreated), whereas dabrafenib treatment of the V600E/T529M mutant increased the closed conformational population less effectively than the V600E mutant alone (1.4 fold vs. 2 fold). This again agrees with the Resistor predictions, namely that V600E/T529I remains sensitive to dabrafenib but V600E/T529M is resistant. Resistor predicted that the V600E/T529I and V600E/T529M mutants would be resistant to encorafenib, but the KinCon data indicates that these mutants may actually retain sensitivity to encorafenib, as the inhibitor induces BRAF’s closed state.

In addition, all inhibitors except dabrafenib were predicted to be sensitive against the G466V mutation and showed closing the of kinase conformation [63]. However, in case of dabrafenib, the response was comparable to vemurafenib, although vemurafenib was classified as sensitive. Notably, previous KinCon experiments showed that G466V (and G466R and G466E [66], see below) impaired kinase function consistent with the reduced endogenous ligand binding predicted by Resistor (see “All BRAF Predictions” supplementary table) [63].

In addition to the above retrospective validation, we chose a few mutations and evaluated them using the KinCon reporter. We selected the mutants G466E, G466R, V471F, L505H, and G593D because they were prioritized by Resistor for at least one of the investigated inhibitors and were reported as patient mutations in either the COSMIC [58] or cBioPortal (using the curated set of non-redundant studies) [67, 68] databases (see Table 2).

**Table 2:** Prioritized BRAF mutations selected for experimental testing. We selected these mutants because they were prioritized by Resistor for at least one of the investigated inhibitors and were reported as patient mutations in either the COSMIC or cBioPortal. The numbers in the first four columns indicate the Resistor-predicted Pareto rank with melanoma mutational probabilities. The numbers in the last two columns indicate the number of patient samples containing the mutation reported in the respective database (access date 12/01/2022). Absence of a Pareto rank indicates Resistor predicted the mutant would remain sensitive to the drug.

| Mutation | Vemurafenib | Dabrafenib | Encorafenib | PLX8394 | COSMIC | cBioPortal |
| --- | --- | --- | --- | --- | --- | --- |
| G466E | - | 1 | - | - | 49 | 31 |
| G466R | - | 1 | - | - | 17 | 3 |
| V471F | - | - | 2 | 3 | 5 | 2 |
| L505H | - | - | 3 | - | 8 | 10 |
| G593D | 1 | 1 | 1 | 1 | 4 | 0 |

The expression-normalized basal biosensor signal suggests that both G466E and G466R mutants shift the conformation to an opened state, comparable to the highly oncogenic V600E variant and similar to the effect of the common non-small-cell lung cancer mutation G466V [63]. The V471F, L505H and G593D mutations, in contrast, did not appear to induce a change in the active conformation (Fig. 3B). When exposed to BRAF inhibitors (Fig. 3C), G466E and G466R mutants showed the highest fold increase of the biosensor signal for all four inhibitors tested. The majority of inhibitors, three out of four, were predicted as sensitive against these mutants. Resistor predicted G466E and G466R to be resistant to dabrafenib, and while Resistor predicted dabrafenib had lower sensitivity compared to encorafenib and PLX8394 (which is consistent with the KinCon results), dabrafenib-treated mutants shifted to a closed comformation at least as much as vemurafenib-treated mutants did. The L505H and G593D KinCon mutants were not affected by any inhibitors, as those mutations do not shift the kinase into an active opened kinase conformation which is required for inhibitor binding. While vemurafenib and dabrafenib do not appear to affect the V471F mutant, encorafenib and PLX8394 did induce a closing of the kinase, suggesting that the structural properties of the inhibitor determine the binding affinity to this mutant. This is particularly intriguing, given that the V471F mutation was selected because we predicted it would confer resistance to encorafenib and PLX8394. While the KinCon results suggest that these two compounds still retain binding to the V471F mutant, the mutant itself did not induce a significant opening of the kinase confirmation required for ligand binding. For the latter three mutations (i.e. L505H, G593D, and V471F), it would therefore be required to induce the open conformation some other way, for example by introducing the V600E mutation similar to T529I and T529M described above, to investigate whether resistance would develop to the inhibitors [62].

#### 3.2.2. Retrospective validation of Resistor predictions using BRAF saturation mutagenesis experiments

Wagenaar et al’s 2014 study examined the effects of BRAF inhibitor binding site mutations on inhibitor efficacy [61]. To do so, they carried out targeted saturation mutagenesis on the BRAF vemurafenib binding site in the A375 human melanoma cell line and challenged the mutants with vemurafenib over a three week period [61]. They then sequenced the emergent clones and measured the IC_50_ values of a subset of the mutants. Importantly, they demonstrated correlation between a mutant’s deep sequencing enrichment, i.e. the increase in the amount of an amino acid sequence in a sample before and after the addition of an inhibitor, and its IC_50_ value [61]. We therefore compared their enrichment data to the Resistor predictions and determined Resistor’s vemurafenib resistance prediction specificity to be 91%. There were five Resistor-predicted resistance mutations that had increased enrichment over the three week period: T529M already discussed above (enriched 47.96 fold above the V600E baseline, which was the experiment’s largest change in enrichment), T529L (enriched 18.57 fold above baseline), T529F (enriched 7.87 fold above baseline), G593I (enriched 4.84 fold above baseline), and L514E (enriched 3.73 fold above baseline). Furthermore, Wagenaar et al. determined the relative IC_50_ values of T529M, T529L, and G593I which were, respectively, 2.05, 2.16, and 3.19 times larger than the IC_50_ for vemurafenib applied to the V600E mutant. The IC_50_ of T529F and L514E were not determined.

To further elucidate the molecular mechanisms conferring resistance to the G593I and L514E mutants, we analyzed the OSPREY-predicted structural models. While neither mutant requires a movement of vemurafenib (Fig. 4A) akin to what was observed in the EGFR and osimertinib structures (Fig. 2), the mutations still lead to a loss of favorable interactions and/or the introduction of energetically unfavorable contacts. The residue G593 (Fig. 4B) may facilitate structural adaptions required for BRAF to accommodate the vemurafenib propyl sulfonamide moiety in the rear of the ATP binding site and the G593L mutations may thus constrain the flexibility of this loop region. In addition, the leucine side chain may project near to the fluoro-substituted central phenyl ring and introduce steric clashes (Fig. 4C). The neighboring D594 backbone interacts with the vemurafenib sulphonamide-nitrogen (Fig. 4B), and this interaction would be weakened in the G593L mutant. Furthermore, residue L514 makes a range of hydrophobic contacts with vemurafenib (Fig. 4D), including the central phenyl ring and the propyl-chain, which are lost in the L514E mutant (Fig. 4E).

**Fig. 4:**
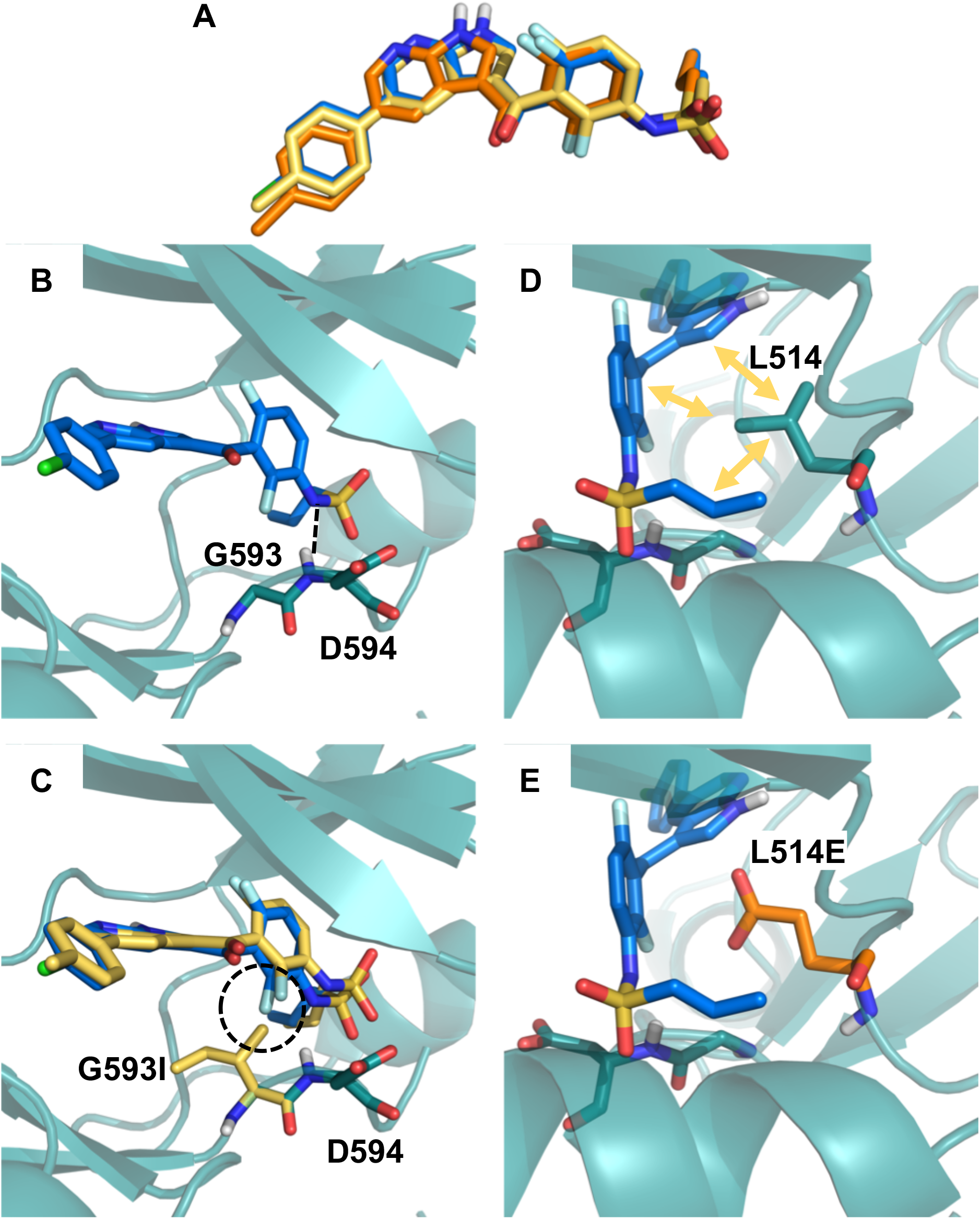
Structural analysis of BRAF mutations G593I and L514E. (A) No major movements were required for vemurafenib to bind to the G593I (yellow) and L514E (orange) mutation in comparison to the wild type binding pose (blue). (B) BRAF G593 is located on the N-terminus of the activation loop and may facilitate conformational changes required to accommodate the vemurafenib propyl sulfonamide moiety in the back of the pocket. The backbone of the neighboring D594 residue interacts with the sulfonamide nitrogen of vemurafenib as indicated by black dashed lines. (C) Mutation of G593 to L not only restricts flexibility of the loop, but also puts the leucine side chain in too close proximity to the fluoro-substituted phenyl ring (highlighted with the dashed circle). (D) Residue L514 is involved in a variety of hydrophobic contacts with vemurafenib (indicated by yellow arrows), which are lost in the L514E mutant (E).

#### 3.3. Complexity

There are a number of distinct steps in Resistor, each of which has its own complexity. While there are sublinear *K*^*^ algorithms, such as BBK^*^ [69] with MARK^*^ [49], these algorithms so far have only been applied to positive and negative design with optimization of specific multiple objectives, such as minimizing/maximizing the bound (respectively unbound) state partition functions and their ratios for computing binding affinity or stability. Comets [32] provably does multistate design optimizing arbitrary constrained linear combinations of GMEC energies, but Comets does not model the partition functions required for calculating binding affinity. A provable ensemble-based algorithm analogous to Comets for arbitrary multistate design optimization is yet to be developed. Thus, general multistate *K*^*^ design remains, unfortunately, a problem linear in the number of sequences and thus exponential in the number of mutable residues.

Computing *K*^*^ itself, as a ratio of partition functions built from the thermodynamic ensembles of the bound to unbound states, can be expensive [70–72]. In order to reduce the number of *K*^*^ problems to solve, Comets is employed as a pruning mechanism for all sequences in which there are more than one mutation. Without Comets, Resistor would need to compute *sN K*^*^ scores, where *s* is the number of states and *N* is the number of sequences. With Comets, Resistor is able to avoid computing many of these *K*^*^ scores, as Comets has been shown in practice to reduce the number of required GMEC calculations by over 99% and to reduce *N* for continuous designs by 96%, yielding an overall speedup of over 5 × 10^5^-fold [32]. Since in this study we considered only single residue mutations we omitted the Comets pruning step, but in any use of Resistor that considers multiple simultaneously mutable residues we believe Comets’ empirical sublinearity will make the difference between feasible and infeasible searches.

Moreover, by using an approximation containing fixed partition function size and sparse residue interaction graphs, we can use the BWM* algorithm [73] to compute the *K*\* scores in time 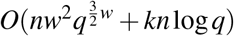, where *w* is the branch-width and *q* the number of rotamers per residue. When we have *w* = *O*(1) this is polynomial time. In this study we found that the *ε*-approximation algorithms using adaptively-sized partition functions, such as BBK^*^ with MARK^*^, were fast enough. However, for larger problems the sparse approximations allow us to approximate the necessary *K*^*^ scores for resistance prediction in time exponential only in the branch-width, and thus polynomial time for fixed branch-widths.

## 4. Discussion

In this work, we report Resistor, a computational algorithm to systematically investigate protein mutations and identify those that have a high likelihood of lowering drug potency in comparison to native substrates. In addition, we analyze the probability that such a mutation is generated in cancer patients and thus likely of clinical importance. Our algorithm is novel, bringing the power of Pareto optimization and computational protein design together and applying them for the first time to predict resistance mutations. The Pareto ranking provides an objective way of prioritizing the most relevant mutations for experimental testing. In addition, we used computationally predicted input structures of ligand-target complexes whenever experimental data was lacking. This is an important step towards expanding the applicability of Resistor, as we have found that, perhaps surprisingly, the availability of high-resolution experimental ligand-target structures still can present a major bottleneck in computational protein design.

We have applied Resistor to two case studies, EGFR and BRAF, in a retrospective manner and, in case of BRAF, also included prospective experimental data for validation. In EGFR and BRAF, the algorithm correctly identified resistance mutations. Using the vemurafenib data by Wagenaar and colleagues [61], which is the most comprehensive dataset on BRAF mutations and vemurafenib resistance, we determined RESISTOR’s vemurafenib resistance prediction specificity and sensitivity to be 91% and 31%, respectively. In a data-rich setting such as proteomics (e.g. [74]), the sensitivity could be regarded as low. However, the prediction of antineoplastic resistance mutations is a sparse data problem. Comprehensive datasets on drug resistance mutations on specific targets are virtually non-existent. We speculate that the reason for this can be found in the large number of individual mutants that must be generated and tested. For example, in our study we used Resistor to investigate 462, 438, and 357 individual mutants for Erlotinib, Gefitinib, and Osimertinib, respectively. While this is computationally feasible, it far exceeds the testing capacities of most experimental groups. Clinical resistance data is even more limited. Furthermore, even for those mutations that have been confirmed to confer clinical resistance in patients, the underlying molecular mechanisms often remain uninvestigated.

Resistor prioritizes escape mutations causing ablation of inhibitor binding and/or tighter substrate binding (the latter as a proxy for *K*_M_). However, mutations affecting the drug target could also mediate resistance via other molecular processes such as altering the stability of conformational states or protein-protein interactions. In addition, clinical resistance is caused by several different mechanisms, of which the relative importance of escape mutations can vary greatly. In some kinases, such as c-Abl, EGFR, and FLT3, active site escape mutations are the main cause of acquired resistance [75]. In other kinases, such as BRAF, escape mutations are not the main mechanism of acquired resistance [76]. Rather, splice variants, amplification, and mutations in related genes such as N-RAS, MEK1, MEK2, IGF-1R, and AKT comprise the majority of cases of clinical resistance [76]. From this perspective, the specificity of Resistor for BRAF and vemurafenib is remarkable and the sensitivity is in line with the fraction of resistance mutations whose aetiology is definitively escape via active site mutation.

We believe that the remaining gap can be closed in future work by modelling additional conformational flexibility, kinetics, and the protein-protein interactions of additional effectors. Yet, despite these limitations, Resistor is able to prioritize mutations that are demonstrated to confer resistance in patients. Specifically, our results show that detailed and combinatorial thermodynamic computations can form the basis for predicting escape mutations to TKIs. In the future, since some resistance mutations exploit kinetic phenomena, kinetics could be incorporated for a more comprehensive model.

## 5. Conclusions

Resistor fills an important void in the science of predicting resistance mutations by providing an algorithm to enumerate the entire Pareto frontier of multiple resistance-causing criteria. It is, to our knowledge, the first algorithm to apply Pareto optimization to predicting resistance mutations. By categorizing predicted resistance mutations by their Pareto rank, it allows the drug discovery community to prioritize escape mutations on the Pareto frontier. Resistor also provides structural justification for the mechanism of each predicted escape mutation by generating an ensemble of predicted structural models upon mutation. In this study, we have applied Resistor to predict resistance mutations in EGFR and BRAF for a number of different therapeutics. We demonstrate that Resistor can also be applied to computationally generated input structures, although the accuracy of the results may be somewhat diminished compared to experimentally determined structures of target-ligand complexes. However, computationally-derived models can still provide useful insights, especially when considering that the availability of experimental structures appears as major bottleneck. While Resistor as described herein optimizes over 4 objectives, as a general method any number of diverse objectives could be added. Resistor can be applied not only to cancer therapeutics, but also to antimicrobial or antiviral drug design. Thus it provides an important tool to the drug discovery community to design drugs that are less prone to resistance.

## Acknowledgements

BRD & NG: We thank all members of the Donald lab for helpful discussions and the NIH (grants R01-GM078031, R01-GM118543, and R35-GM144042 to BRD) for funding. TK, ES, & AF were funded in whole, or in part, by the Austrian Science Fund (FWF) P34376, P27606, P30441, P32960, and P35159. For the purpose of open access, the author has applied a CC-BY public copyright license to any Author Accepted Manuscript version arising from this submission.

## Competing interests

BRD is a founder of Ten63 Therapeutics, Inc.

### Box 1

Progress and Potential

Targeted cancer drugs developed over the past two decades have been instrumental in treating certain types of cancer and extending patient lifespans. These drugs include kinase inhibitors targeting EGFR and BRAF, two important enzymes of the mitogen-activated protein kinase pathway whose dysregulation can lead to many types of cancer, including melanoma and non-small cell lung cancer. The inhibitors are effective for a period of time but the tumors often develop resistance to the drugs, leading once again to cancer progression. The ability to predict how an enzyme target can develop drug resistance would allow for a proactive, resistance-aware approach to drug design. Our new algorithm Resistor uses structure-based computational design to predict how different mutations in an enzyme will affect a drug’s efficacy. It pairs these predictions with empirical data on how likely a mutation is to occur in a given cancer type, which allows researchers to identify “mutational hotspots,” or particular places where mutations are most likely to cause drug resistance. These predictions provide designers new insights during the drug development process that should allow for the quicker development of more durable and longer-lasting cancer therapeutics.

## STAR Methods

### Resource Availability

#### Lead contact

Further information and requests for resources and reagents should be directed to and will be fulfilled by the lead contact, Bruce Donald.

#### Materials availability

Materials are available upon request to the Lead Contact.

#### Data and code availability

- OSPREY design specifications and mutational signature probabilities required to reproduce the predictions in this paper have been deposited at in the Harvard Dataverse and are publicly available as of the date of publication. DOIs are listed in the key resources table.
- The version of OSPREY used in this paper has been deposited in the Harvard Dataverse and is publicly available as of the date of publication. DOIs are listed in the key resources table. For new empirical designs, we recommend using the latest version of OSPREY available for free at http://www.cs.duke.edu/donaldlab/osprey.php. All code for the OSPREY software package is also available on GitHub at https://github.com/donaldlab/OSPREY3, and is free and open-source.
- Any additional information required to reanalyze the data reported in this paper is available from the lead contact upon request.

### Experimental Model and Subject Details

#### Cell Culture and Antibodies

HEK293T cells were grown in Dulbecco’s Modified Eagle Medium (DMEM) supplemented with 10% fetal bovine serum (FBS). Transient transfections were performed with Transfectin reagent (Bio-Rad, 1703352). Mouse anti-BRAF (Santa Cruz, F-7: sc-5284) antibody was used to determine biosensor expression levels.

### Method Details

#### Preparation of Empirical and Docked Structures for *K*^*^ Predictions

The crystal structures used for the EGFR predictions were adopted from Kaserer and Blagg’s 2018 publication [20]. A full description of the PDB entries used can be found in that paper’s section *Table S7*, and details on how the structures were prepared for OSPREY predictions is in that paper’s section *Structure Selection and Preparation*.

For BRAF, the crystal structures of vemurafenib (PDB id 3og7 [48]) and dabrafenib (PDB id 4xv2 [47]) in complex with BRAF V600E were selected as input for Resistor. Both structures have been prepared using the default setting of the Protein Preparation Wizard [77] in Maestro [78]. In the case of encorafenib and PLX8394, crystal structures of structurally closely related, but not the identical, molecules were available. These experimental complexes were used to generate encorafenib and PLX8394 models. Encorafenib was docked into PDB id 4xv3 [47] using the default settings of the induced fit docking procedure in Maestro [78–81]. For validation, the co-crystallized ligand PLX7922 was re-docked. The highest scored docking pose of encorafenib was selected for further investigation. We found that the conserved substructures in encorafenib and PLX7922 aligned very well in this docking pose.

For PLX8394, re-docking of the co-crystallized ligand PLX7904 (PDB id 4xv1 [47]) failed with the induced fit docking procedure, but was successful using a rigid docking workflow in GOLD version 5.8.0 [82]. The binding site was defined as 6 Å around the ligand and the water molecule HOH905 was set to toggle and spin. The default settings of all other parameters were used.

An experimental structure of the endogenous ligand ADP was available, however, BRAF adopted in inactive conformation in this complex. Apo BRAF in its active conformation (PDB id 4mne [83]) was thus combined with ANP-bound protein kinase c-src (PDB id 2src [84]) to generate an active, endogenous ligand-bound BRAF complex. This model was used as template to build a BRAF:ADP homology model in the Molecular Operating Environment [85] using the default settings. This included refinement steps to resolve potential steric clashes in the rather crude ANP-BRAF input template.

For all complexes, water molecules not involved in mediating interactions between the ligand and the target were deleted and only residues with a 12 Å radius around the ligand were kept in the final input structures.

#### Evaluation of Ligand Affinity

The command line interface of OSPREY was used to generate distinct YAML design files for each residue within 5 Å of a ligand. These YAML design files specify the input structures, the mutable residues, the flexible residues, and connectivity templates for OSPREY. To create the forcefield parameters files for the inhibitors and endogenous ligands, we used the Antechamber program in the AmberTools software package [86]. Then, to calculate the *K*^*^ scores we used OSPREY with the following command input:

~~~
osprey affinity --design <YAML design file> --epsilon 0.63 --frcmod <force field modification file> --stability-threshold -1
~~~

where `<YAML design file>` was replaced with the individual YAML design file and `<force field modification file>` was replaced with the AmberTools-generated file. The YAML design and forcefield modification files used in this study are available in the Harvard Dataverse (see Key Resources Table).

#### Luciferase PCA analyses

We transiently overexpressed indicated versions of the Rluc-PCA–based KinCon biosensors in 24-well plate formats. Experiments were performed 48h post transfection. For the luciferase-PCA measurements, the growth medium was carefully removed and the cells were washed with phosphate-buffered saline (PBS). Cell suspensions were transferred to 96-well plates and subjected to luminescence analysis using the PHERAstar FSX (BMG Labtech). Luciferase luminescence signals were integrated for 10 seconds following addition of the Rluc substrate benzyl-coelenterazine (NanoLight, #301). Cell lysates were prepared post RLU measurements. Expression levels of the biosensor were determined via western blot analysis.

### Quantification and Statistical Analysis

In Fig. 3, the student’s T-test was used to evaluate whether the mean of the RLU of a mutant was significantly different from that of the relative DMSO control. The SEM was used with n = 4. Significance was defined to three different p-levels, where \**p <* 0.05, \*\**p <* 0.01, and \*\*\**p <* 0.001.

### S1. The *K*^*^ algorithm

*K*^*^ is an *ε*-accurate algorithm for computing a provable approximation to the affinity constant *K*_*a*_. It is implemented in the OSPREY computational protein design software package [18, 87]. *K*^*^ is defined as the quotient of the bound to unbound partition functions of a protein:ligand system for a given amino acid sequence. For a proof that *K*^*^ approximates *K*_*a*_ see Appendix A of [87].

*K*^*^ calculates an *ε*-accurate partition function for three structures: the bound protein:ligand complex (denoted *PL*), the unbound protein (denoted *P*), and the unbound ligand (denoted *L*). Let *X* be an arbitrary state, *X* ∈ {*P, L, PL*}. The partition function is a summation of the Boltzmann-weighted energies for all of the conformations in the thermodynamic ensemble of *X*. Let *s* denote an arbitrary amino acid sequence, then the partition function of *s* in state *X* (which we donate as *q*_*X*_ (*s*)) is:

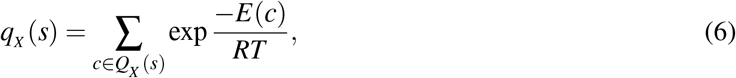

where *Q*_*X*_ (*s*) is the entire conformational ensemble of sequence *s* in state *X*, and *c* is a single conformation from that ensemble. *E*(*c*) is the energy of conformation *c. R* is the ideal gas constant and *T* is the temperature in absolute Kelvin.

The *K*^*^ score for a sequence *s* approximates *K*_*a*_:

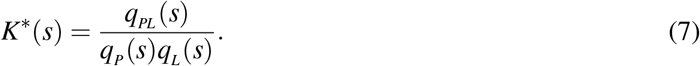

By using an *A*^*^ search over *Q*_*X*_ (*s*) to generate an ordered, gap-free list of low energy conformations, the *K*^*^ algorithms generates an *ε*-approximation of the partition function *q*_*X*_ (*s*) and the ensemble-complete *K*^*^ value. This approximation is known as the *K*^*^ score.

Inputs to the *K*^*^ algorithm include 1) an input structure; 2) a conformation library; 3) an energy function; 4) *ε*, and; 5) flexibility and mutability choices.

### S2. Empirical Resistor runtimes

The Resistor computation entails three stages: 1) computing the positive and negative *K*^*^ designs; 2) assigning mutational signature probabilities to each mutation, and; 3) run Pareto optimization over the four axes. Steps 2 and 3 empirically take a negligible amount of time, on the order of seconds. Step 1, however, computes two partition functions for each sequence and can take more time. Figure S1 shows the empirical runtime (in seconds) that it took our computers to run the positive and negative *K*^*^ designs, where a design mutated a residue to each of the 19 other possible amino acids.

**Fig. S1:**
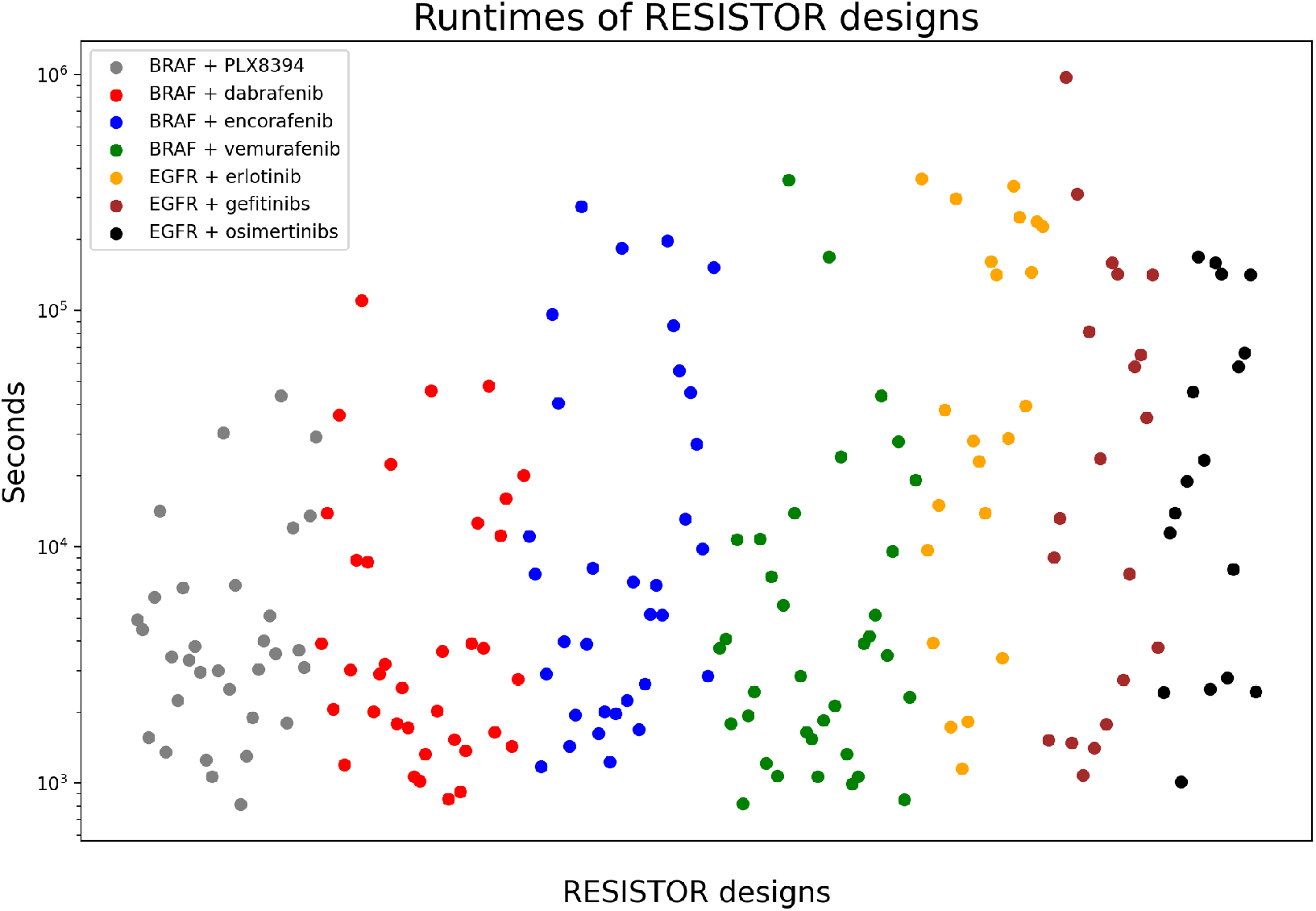
Positive and negative design runtimes for a residue location. Each dot represents the amount of time (in seconds) that Resistor took to compute the positive and negative *K*^*^ designs for a given mutation location. This means that each dot represents the computation of 40 *K*^*^ scores. The computation times range from 813 seconds to 972465 seconds, with the average being 40630 seconds or 1015 seconds per sequence. The different colors represent the particular kinase/inhibitor pair. The designs were run on a 24-core, 48-thread Intel Xeon processor with 4 Nvidia Titan V GPUs.

### S3. EGFR Pareto Frontier

**Table S1:**
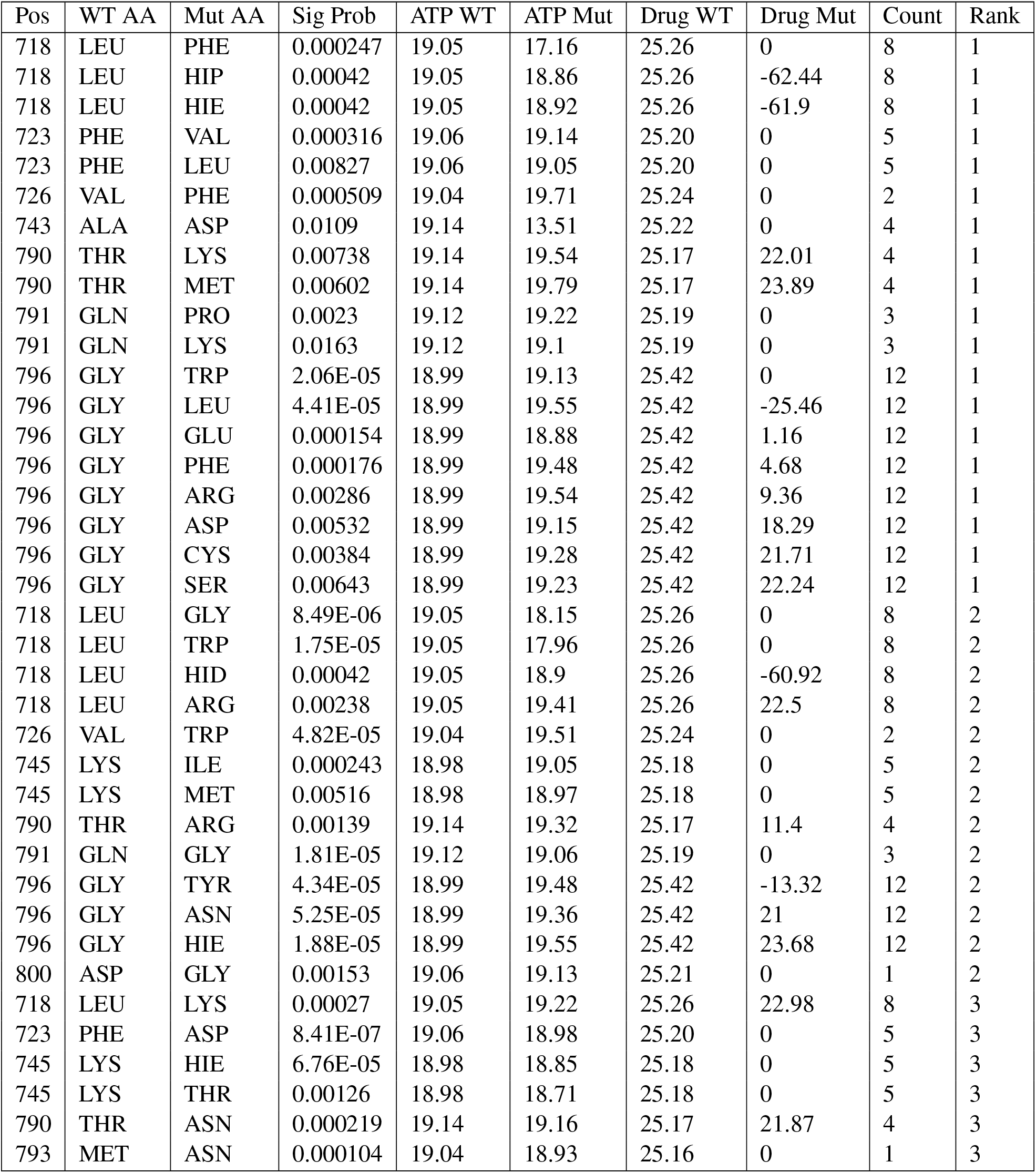
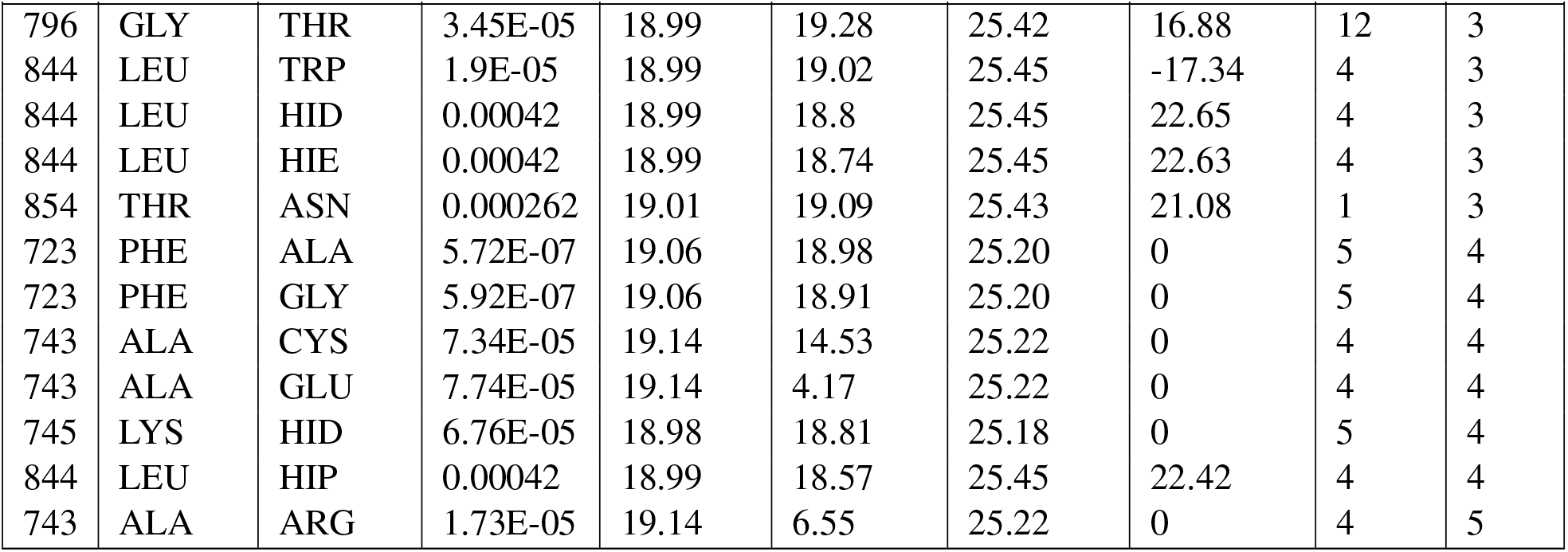
All Resistor resistance mutation predictions for EGFR with erlotinib. “Pos” is the position of the residue. “WT AA” is the wildtype identity of the amino acid. “Mut AA” is the resistance mutation. “Sig Prob” is the mutational signature probability for the mutation from “WT AA” to “Mut AA” in lung adenocarcinoma. “ATP WT” and “ATP Mut” are the *K*^*^ scores of the endogenous ligand with the wildtype and mutant residues, respectively. “Drug WT” and “Drug Mut” are the *K*^*^ scores of erlotinib with the wildtype and mutant residues, respectively. “Count” is number of resistance mutations at the position. “Rank” is the Pareto rank of the mutation. Note: *K*^*^ scores are in log_10_ units where possible and 0 where there is predicted to be no binding.

| Pos | WT AA | Mut AA | Sig Prob | ATP WT | ATP Mut | Drug WT | Drug Mut | Count | Rank |
| --- | --- | --- | --- | --- | --- | --- | --- | --- | --- |
| 718 | LEU | PHE | 0.000247 | 19.05 | 17.16 | 25.26 | 0 | 8 | 1 |
| 718 | LEU | HIP | 0.00042 | 19.05 | 18.86 | 25.26 | -62.44 | 8 | 1 |
| 718 | LEU | HIE | 0.00042 | 19.05 | 18.92 | 25.26 | -61.9 | 8 | 1 |
| 723 | PHE | VAL | 0.000316 | 19.06 | 19.14 | 25.20 | 0 | 5 | 1 |
| 723 | PHE | LEU | 0.00827 | 19.06 | 19.05 | 25.20 | 0 | 5 | 1 |
| 726 | VAL | PHE | 0.000509 | 19.04 | 19.71 | 25.24 | 0 | 2 | 1 |
| 743 | ALA | ASP | 0.0109 | 19.14 | 13.51 | 25.22 | 0 | 4 | 1 |
| 790 | THR | LYS | 0.00738 | 19.14 | 19.54 | 25.17 | 22.01 | 4 | 1 |
| 790 | THR | MET | 0.00602 | 19.14 | 19.79 | 25.17 | 23.89 | 4 | 1 |
| 791 | GLN | PRO | 0.0023 | 19.12 | 19.22 | 25.19 | 0 | 3 | 1 |
| 791 | GLN | LYS | 0.0163 | 19.12 | 19.1 | 25.19 | 0 | 3 | 1 |
| 796 | GLY | TRP | 2.06E-05 | 18.99 | 19.13 | 25.42 | 0 | 12 | 1 |
| 796 | GLY | LEU | 4.41E-05 | 18.99 | 19.55 | 25.42 | -25.46 | 12 | 1 |
| 796 | GLY | GLU | 0.000154 | 18.99 | 18.88 | 25.42 | 1.16 | 12 | 1 |
| 796 | GLY | PHE | 0.000176 | 18.99 | 19.48 | 25.42 | 4.68 | 12 | 1 |
| 796 | GLY | ARG | 0.00286 | 18.99 | 19.54 | 25.42 | 9.36 | 12 | 1 |
| 796 | GLY | ASP | 0.00532 | 18.99 | 19.15 | 25.42 | 18.29 | 12 | 1 |
| 796 | GLY | CYS | 0.00384 | 18.99 | 19.28 | 25.42 | 21.71 | 12 | 1 |
| 796 | GLY | SER | 0.00643 | 18.99 | 19.23 | 25.42 | 22.24 | 12 | 1 |
| 718 | LEU | GLY | 8.49E-06 | 19.05 | 18.15 | 25.26 | 0 | 8 | 2 |
| 718 | LEU | TRP | 1.75E-05 | 19.05 | 17.96 | 25.26 | 0 | 8 | 2 |
| 718 | LEU | HID | 0.00042 | 19.05 | 18.9 | 25.26 | -60.92 | 8 | 2 |
| 718 | LEU | ARG | 0.00238 | 19.05 | 19.41 | 25.26 | 22.5 | 8 | 2 |
| 726 | VAL | TRP | 4.82E-05 | 19.04 | 19.51 | 25.24 | 0 | 2 | 2 |
| 745 | LYS | ILE | 0.000243 | 18.98 | 19.05 | 25.18 | 0 | 5 | 2 |
| 745 | LYS | MET | 0.00516 | 18.98 | 18.97 | 25.18 | 0 | 5 | 2 |
| 790 | THR | ARG | 0.00139 | 19.14 | 19.32 | 25.17 | 11.4 | 4 | 2 |
| 791 | GLN | GLY | 1.81E-05 | 19.12 | 19.06 | 25.19 | 0 | 3 | 2 |
| 796 | GLY | TYR | 4.34E-05 | 18.99 | 19.48 | 25.42 | -13.32 | 12 | 2 |
| 796 | GLY | ASN | 5.25E-05 | 18.99 | 19.36 | 25.42 | 21 | 12 | 2 |
| 796 | GLY | HIE | 1.88E-05 | 18.99 | 19.55 | 25.42 | 23.68 | 12 | 2 |
| 800 | ASP | GLY | 0.00153 | 19.06 | 19.13 | 25.21 | 0 | 1 | 2 |
| 718 | LEU | LYS | 0.00027 | 19.05 | 19.22 | 25.26 | 22.98 | 8 | 3 |
| 723 | PHE | ASP | 8.41E-07 | 19.06 | 18.98 | 25.20 | 0 | 5 | 3 |
| 745 | LYS | HIE | 6.76E-05 | 18.98 | 18.85 | 25.18 | 0 | 5 | 3 |
| 745 | LYS | THR | 0.00126 | 18.98 | 18.71 | 25.18 | 0 | 5 | 3 |
| 790 | THR | ASN | 0.000219 | 19.14 | 19.16 | 25.17 | 21.87 | 4 | 3 |
| 793 | MET | ASN | 0.000104 | 19.04 | 18.93 | 25.16 | 0 | 1 | 3 |
| 796 | GLY | THR | 3.45E-05 | 18.99 | 19.28 | 25.42 | 16.88 | 12 | 3 |
| 844 | LEU | TRP | 1.9E-05 | 18.99 | 19.02 | 25.45 | -17.34 | 4 | 3 |
| 844 | LEU | HID | 0.00042 | 18.99 | 18.8 | 25.45 | 22.65 | 4 | 3 |
| 844 | LEU | HIE | 0.00042 | 18.99 | 18.74 | 25.45 | 22.63 | 4 | 3 |
| 854 | THR | ASN | 0.000262 | 19.01 | 19.09 | 25.43 | 21.08 | 1 | 3 |
| 723 | PHE | ALA | 5.72E-07 | 19.06 | 18.98 | 25.20 | 0 | 5 | 4 |
| 723 | PHE | GLY | 5.92E-07 | 19.06 | 18.91 | 25.20 | 0 | 5 | 4 |
| 743 | ALA | CYS | 7.34E-05 | 19.14 | 14.53 | 25.22 | 0 | 4 | 4 |
| 743 | ALA | GLU | 7.74E-05 | 19.14 | 4.17 | 25.22 | 0 | 4 | 4 |
| 745 | LYS | HID | 6.76E-05 | 18.98 | 18.81 | 25.18 | 0 | 5 | 4 |
| 844 | LEU | HIP | 0.00042 | 18.99 | 18.57 | 25.45 | 22.42 | 4 | 4 |
| 743 | ALA | ARG | 1.73E-05 | 19.14 | 6.55 | 25.22 | 0 | 4 | 5 |

**Table S2:**
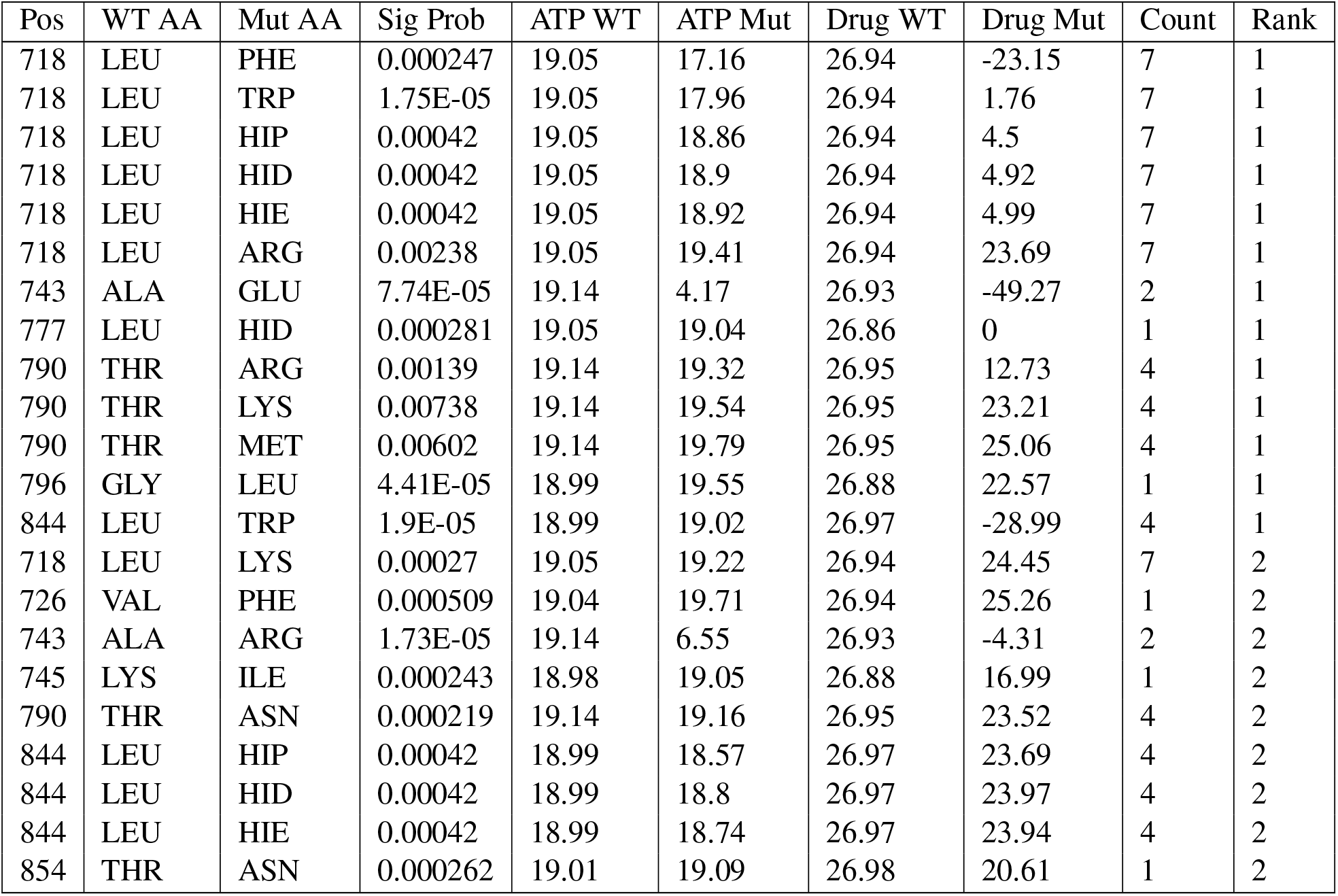
All Resistor resistance mutation predictions for EGFR with gefitinib. “Pos” is the position of the residue. “WT AA” is the wildtype identity of the amino acid. “Mut AA” is the resistance mutation. “Sig Prob” is the mutational signature probability for the mutation from “WT AA” to “Mut AA” in lung adenocarcinoma. “ATP WT” and “ATP Mut” are the *K*^*^ scores of the endogenous ligand with the wildtype and mutant residues, respectively. “Drug WT” and “Drug Mut” are the *K*^*^ scores of gefitinib with the wildtype and mutant residues, respectively. “Count” is number of resistance mutations at the position. “Rank” is the Pareto rank of the mutation. Note: *K*^*^ scores are in log_10_ units where possible and 0 where there is predicted to be no binding.

**Table S3:** All Resistor resistance mutation predictions for EGFR with osimertinib. “Pos” is the position of the residue. “WT AA” is the wildtype identity of the amino acid. “Mut AA” is the resistance mutation. “Sig Prob” is the mutational signature probability for the mutation from “WT AA” to “Mut AA” in lung adenocarcinoma. “ATP WT” and “ATP Mut” are the *K*^*^ scores of the endogenous ligand with the wildtype and mutant residues, respectively. “Drug WT” and “Drug Mut” are the *K*^*^ scores of osimertinib with the wildtype and mutant residues, respectively. “Count” is number of resistance mutations at the position. “Rank” is the Pareto rank of the mutation. Note: *K*^*^ scores are in log_10_ units where possible and 0 where there is predicted to be no binding.

| Pos | WT AA | Mut AA | Sig Prob | ATP WT | ATP Mut | Drug WT | Drug Mut | Count | Rank |
| --- | --- | --- | --- | --- | --- | --- | --- | --- | --- |
| 718 | LEU | TRP | 1.75E-05 | 19.05 | 17.96 | 27.51 | 0 | 9 | 1 |
| 718 | LEU | PHE | 0.000247 | 19.05 | 17.16 | 27.51 | 0 | 9 | 1 |
| 718 | LEU | HIP | 0.00042 | 19.05 | 18.86 | 27.51 | -38.77 | 9 | 1 |
| 718 | LEU | HIE | 0.00042 | 19.05 | 18.92 | 27.51 | -38.57 | 9 | 1 |
| 718 | LEU | MET | 0.0108 | 19.05 | 19.44 | 27.51 | 25.52 | 9 | 1 |
| 719 | GLY | VAL | 0.017 | 19.05 | 14.7 | 27.50 | 20.75 | 2 | 1 |
| 726 | VAL | TRP | 4.82E-05 | 19.04 | 19.51 | 27.48 | 0 | 3 | 1 |
| 743 | ALA | ASP | 0.0109 | 19.14 | 13.51 | 27.43 | 17.5 | 3 | 1 |
| 796 | GLY | TRP | 2.06E-05 | 18.99 | 19.13 | 27.38 | -97.39 | 14 | 1 |
| 796 | GLY | TYR | 4.34E-05 | 18.99 | 19.48 | 27.38 | -61.59 | 14 | 1 |
| 796 | GLY | PHE | 0.000176 | 18.99 | 19.48 | 27.38 | -41.78 | 14 | 1 |
| 796 | GLY | LEU | 4.41E-05 | 18.99 | 19.55 | 27.38 | -3.75 | 14 | 1 |
| 796 | GLY | ARG | 0.00286 | 18.99 | 19.54 | 27.38 | 10.61 | 14 | 1 |
| 796 | GLY | ASP | 0.00532 | 18.99 | 19.15 | 27.38 | 14.96 | 14 | 1 |
| 796 | GLY | CYS | 0.00384 | 18.99 | 19.28 | 27.38 | 21.85 | 14 | 1 |
| 796 | GLY | SER | 0.00643 | 18.99 | 19.23 | 27.38 | 24.76 | 14 | 1 |
| 718 | LEU | HID | 0.00042 | 19.05 | 18.9 | 27.51 | -37.61 | 9 | 2 |
| 718 | LEU | ARG | 0.00238 | 19.05 | 19.41 | 27.51 | 20.16 | 9 | 2 |
| 723 | PHE | ILE | 0.00209 | 19.06 | 19.5 | 27.45 | 25.68 | 1 | 2 |
| 792 | LEU | HIP | 0.00486 | 19.03 | 18.88 | 27.47 | 24.98 | 3 | 2 |
| 792 | LEU | HIE | 0.00486 | 19.03 | 18.93 | 27.47 | 25.07 | 3 | 2 |
| 792 | LEU | HID | 0.00486 | 19.03 | 18.98 | 27.47 | 25.15 | 3 | 2 |
| 796 | GLY | GLU | 0.000154 | 18.99 | 18.88 | 27.38 | 2.49 | 14 | 2 |
| 796 | GLY | HIE | 1.88E-05 | 18.99 | 19.55 | 27.38 | 11.32 | 14 | 2 |
| 796 | GLY | ASN | 5.25E-05 | 18.99 | 19.36 | 27.38 | 18.38 | 14 | 2 |
| 718 | LEU | LYS | 0.00027 | 19.05 | 19.22 | 27.51 | 24.7 | 9 | 3 |
| 719 | GLY | THR | 4.02E-05 | 19.05 | 17.76 | 27.50 | 20.4 | 2 | 3 |
| 726 | VAL | ARG | 2.32E-05 | 19.04 | 17.68 | 27.48 | 21.62 | 3 | 3 |
| 726 | VAL | LYS | 5.39E-05 | 19.04 | 17.07 | 27.48 | 21.87 | 3 | 3 |
| 743 | ALA | GLU | 7.74E-05 | 19.14 | 4.17 | 27.43 | 0.47 | 3 | 3 |
| 796 | GLY | HID | 1.88E-05 | 18.99 | 19.49 | 27.38 | 11.69 | 14 | 3 |
| 796 | GLY | THR | 3.45E-05 | 18.99 | 19.28 | 27.38 | 22.43 | 14 | 3 |
| 844 | LEU | TRP | 1.9E-05 | 18.99 | 19.02 | 27.56 | 22.12 | 1 | 3 |
| 796 | GLY | HIP | 1.88E-05 | 18.99 | 19.46 | 27.38 | 12.09 | 14 | 4 |
| 718 | LEU | GLY | 8.49E-06 | 19.05 | 18.15 | 27.51 | 24.02 | 9 | 5 |
| 743 | ALA | ARG | 1.73E-05 | 19.14 | 6.55 | 27.43 | 12.58 | 3 | 5 |

### S4. BRAF Pareto Frontier

**Table S4:** All Resistor resistance mutation predictions for BRAF with dabrafenib. “Pos” is the position of the residue. “WT AA” is the wildtype identity of the amino acid. “Mut AA” is the resistance mutation. “Sig Prob” is the mutational signature probability for the mutation from “WT AA” to “Mut AA” in melanoma. “ATP WT” and “ATP Mut” are the *K*^*^ scores of the endogenous ligand with the wildtype and mutant residues, respectively. “Drug WT” and “Drug Mut” are the *K*^*^ scores of dabrafenib with the wildtype and mutant residues, respectively. “Count” is number of resistance mutations at the position. “Rank” is the Pareto rank of the mutation. Note: *K*^*^ scores are in log_10_ units where possible and 0 where there is predicted to be no binding.

| Pos | WT AA | Mut AA | Sig Prob | ATP WT | ATP Mut | Drug WT | Drug Mut | Count | Rank |
| --- | --- | --- | --- | --- | --- | --- | --- | --- | --- |
| 466 | GLY | ARG | 5.84E-02 | 18.80 | 10.53 | 37.16 | -167.92 | 11 | 1 |
| 466 | GLY | LYS | 2.38E-02 | 18.80 | 11.79 | 37.16 | -52.4 | 11 | 1 |
| 466 | GLY | GLU | 2.19E-01 | 18.80 | 12.89 | 37.16 | 21.32 | 11 | 1 |
| 471 | VAL | LEU | 4.43E-04 | 18.65 | 19.67 | 37.26 | 25.1 | 6 | 1 |
| 508 | THR | ARG | 2.95E-04 | 18.59 | 18.59 | 37.23 | -118.81 | 4 | 1 |
| 535 | SER | PRO | 1.30E-03 | 18.72 | 18.65 | 37.26 | 0 | 1 | 1 |
| 593 | GLY | PHE | 1.99E-06 | 18.66 | 20.07 | 37.17 | 0 | 16 | 1 |
| 593 | GLY | TYR | 3.58E-05 | 18.66 | 19.86 | 37.17 | 0 | 16 | 1 |
| 593 | GLY | ARG | 7.80E-04 | 18.66 | 16.17 | 37.17 | 0 | 16 | 1 |
| 593 | GLY | GLU | 2.76E-04 | 18.66 | 18.73 | 37.17 | -60.32 | 16 | 1 |
| 593 | GLY | ASN | 1.34E-03 | 18.66 | 19.16 | 37.17 | -39.82 | 16 | 1 |
| 593 | GLY | ASP | 1.63E-02 | 18.66 | 18.89 | 37.17 | -29.79 | 16 | 1 |
| 593 | GLY | CYS | 1.66E-03 | 18.66 | 19.06 | 37.17 | 17.46 | 16 | 1 |
| 593 | GLY | VAL | 9.35E-04 | 18.66 | 19.18 | 37.17 | 28.45 | 16 | 1 |
| 593 | GLY | ILE | 4.55E-05 | 18.66 | 19.8 | 37.17 | 30.24 | 16 | 1 |
| 593 | GLY | SER | 6.09E-02 | 18.66 | 18.83 | 37.17 | 34.27 | 16 | 1 |
| 466 | GLY | GLN | 7.24E-05 | 18.80 | 12.55 | 37.16 | 11.37 | 11 | 2 |
| 466 | GLY | ASP | 7.51E-04 | 18.80 | 17.06 | 37.16 | 18.64 | 11 | 2 |
| 466 | GLY | VAL | 2.51E-03 | 18.80 | 13.44 | 37.16 | 29.23 | 11 | 2 |
| 467 | SER | PRO | 7.37E-04 | 18.62 | 18.85 | 37.15 | 30.42 | 1 | 2 |
| 481 | ALA | LYS | 2.74E-04 | 18.58 | 17.32 | 36.98 | -4.22 | 8 | 2 |
| 481 | ALA | LEU | 7.06E-05 | 18.58 | 18.68 | 36.98 | 9.97 | 8 | 2 |
| 481 | ALA | GLU | 1.21E-03 | 18.58 | 17.92 | 36.98 | 22.11 | 8 | 2 |
| 505 | LEU | ARG | 8.61E-04 | 18.59 | 18.58 | 36.85 | 16.53 | 5 | 2 |
| 508 | THR | LYS | 9.16E-04 | 18.59 | 18.59 | 37.23 | 27.22 | 4 | 2 |
| 514 | LEU | ARG | 5.22E-05 | 18.57 | 17.18 | 37.10 | 21.32 | 12 | 2 |
| 514 | LEU | ILE | 1.65E-03 | 18.57 | 18.4 | 37.10 | 32.55 | 12 | 2 |
| 529 | THR | PHE | 3.77E-05 | 18.58 | 15.91 | 36.99 | -125.31 | 11 | 2 |
| 529 | THR | MET | 1.74E-05 | 18.58 | 18.65 | 36.99 | -8.16 | 11 | 2 |
| 529 | THR | ASN | 9.96E-04 | 18.58 | 18.55 | 36.99 | 34.54 | 11 | 2 |
| 593 | GLY | HIE | 1.66E-05 | 18.66 | 19.58 | 37.17 | 0 | 16 | 2 |
| 593 | GLY | THR | 2.67E-05 | 18.66 | 19.06 | 37.17 | 27.33 | 16 | 2 |
| 464 | GLY | GLN | 7.24E-05 | 18.58 | 2.95 | 37.09 | 11.05 | 1 | 3 |
| 466 | GLY | THR | 5.47E-05 | 18.80 | 14.85 | 37.16 | 29.21 | 11 | 3 |
| 481 | ALA | ILE | 5.14E-05 | 18.58 | 14.2 | 36.98 | 21.38 | 8 | 3 |
| 481 | ALA | VAL | 1.46E-03 | 18.58 | 17.77 | 36.98 | 33.45 | 8 | 3 |
| 505 | LEU | SER | 2.78E-05 | 18.59 | 18.58 | 36.85 | 35.02 | 5 | 3 |
| 514 | LEU | PRO | 1.21E-03 | 18.57 | 18.12 | 37.10 | 33.34 | 12 | 3 |
| 527 | ILE | LEU | 6.37E-05 | 18.60 | 18.61 | 37.30 | 33.5 | 1 | 3 |
| 593 | GLY | HIP | 1.66E-05 | 18.66 | 19.56 | 37.17 | 0 | 16 | 3 |
| 514 | LEU | SER | 2.27E-04 | 18.57 | 18.1 | 37.10 | 34.19 | 12 | 4 |
| 593 | GLY | HID | 1.66E-05 | 18.66 | 19.55 | 37.17 | 0 | 16 | 4 |
| 471 | VAL | MET | 8.44E-06 | 18.65 | 18.96 | 37.26 | 27.83 | 6 | 5 |
| 481 | ALA | ARG | 1.53E-05 | 18.58 | 17.02 | 36.98 | -15.13 | 8 | 5 |
| 481 | ALA | ASP | 4.39E-06 | 18.58 | 19.21 | 36.98 | 31.9 | 8 | 5 |
| 508 | THR | GLU | 1.05E-05 | 18.59 | 18.59 | 37.23 | 35 | 4 | 5 |
| 514 | LEU | PHE | 1.53E-05 | 18.57 | 18.19 | 37.10 | 0 | 12 | 5 |
| 578 | LYS | TYR | 5.24E-06 | 18.55 | 18.39 | 37.11 | -143.76 | 1 | 5 |
| 593 | GLY | TRP | 4.78E-06 | 18.66 | 18.83 | 37.17 | 0 | 16 | 5 |
| 593 | GLY | LEU | 3.87E-07 | 18.66 | 19.35 | 37.17 | -85.52 | 16 | 5 |
| 466 | GLY | PRO | 1.77E-07 | 18.80 | 18.27 | 37.16 | 0 | 11 | 6 |
| 466 | GLY | TRP | 2.10E-06 | 18.80 | 13.1 | 37.16 | 0 | 11 | 6 |
| 466 | GLY | LEU | 4.82E-06 | 18.80 | 9.74 | 37.16 | -39.41 | 11 | 6 |
| 466 | GLY | CYS | 3.82E-06 | 18.80 | 17.62 | 37.16 | 27.45 | 11 | 6 |
| 469 | GLY | PRO | 1.77E-07 | 18.60 | 18.75 | 37.10 | 0 | 1 | 6 |
| 471 | VAL | PRO | 5.74E-07 | 18.65 | 17.88 | 37.26 | 0 | 6 | 6 |
| 471 | VAL | ARG | 3.03E-07 | 18.65 | 18.99 | 37.26 | 23.05 | 6 | 6 |
| 471 | VAL | GLU | 9.01E-07 | 18.65 | 18.82 | 37.26 | 31.61 | 6 | 6 |
| 481 | ALA | GLN | 1.80E-06 | 18.58 | 18.37 | 36.98 | 15.47 | 8 | 6 |
| 508 | THR | GLN | 2.51E-06 | 18.59 | 18.59 | 37.23 | 33.59 | 4 | 6 |
| 513 | ILE | ARG | 1.06E-06 | 18.57 | 18.57 | 37.09 | -30.17 | 2 | 6 |
| 513 | ILE | TYR | 2.73E-06 | 18.57 | 18.57 | 37.09 | 27.86 | 2 | 6 |
| 514 | LEU | HIP | 3.37E-07 | 18.57 | 17.68 | 37.10 | 25.98 | 12 | 6 |
| 514 | LEU | HID | 3.37E-07 | 18.57 | 17.72 | 37.10 | 26.06 | 12 | 6 |
| 514 | LEU | LYS | 1.68E-06 | 18.57 | 17.37 | 37.10 | 32.25 | 12 | 6 |
| 514 | LEU | MET | 2.51E-06 | 18.57 | 18.42 | 37.10 | 33.68 | 12 | 6 |
| 528 | VAL | ARG | 5.57E-07 | 18.57 | 18.61 | 37.01 | -72.34 | 1 | 6 |
| 529 | THR | TYR | 1.12E-06 | 18.58 | 18.58 | 36.99 | -10.9 | 11 | 6 |
| 529 | THR | ARG | 4.98E-07 | 18.58 | 18.6 | 36.99 | -4.95 | 11 | 6 |
| 529 | THR | LYS | 1.74E-06 | 18.58 | 18.57 | 36.99 | 4.7 | 11 | 6 |
| 529 | THR | LEU | 3.10E-06 | 18.58 | 18.56 | 36.99 | 26.71 | 11 | 6 |
| 532 | CYS | HID | 4.44E-06 | 18.49 | 14.39 | 37.09 | 26.54 | 7 | 6 |
| 532 | CYS | HIP | 4.44E-06 | 18.49 | 14.51 | 37.09 | 26.79 | 7 | 6 |
| 532 | CYS | HIE | 4.44E-06 | 18.49 | 12.48 | 37.09 | 25.2 | 7 | 6 |
| 532 | CYS | ILE | 1.16E-06 | 18.49 | 18.82 | 37.09 | 34.28 | 7 | 6 |
| 532 | CYS | VAL | 2.11E-06 | 18.49 | 18.7 | 37.09 | 34.77 | 7 | 6 |
| 471 | VAL | HID | 6.58E-07 | 18.65 | 17.61 | 37.26 | 33.79 | 6 | 7 |
| 505 | LEU | GLY | 6.90E-08 | 18.59 | 18.58 | 36.85 | 34.62 | 5 | 7 |
| 505 | LEU | GLN | 1.84E-06 | 18.59 | 18.6 | 36.85 | 34.82 | 5 | 7 |
| 514 | LEU | HIE | 3.37E-07 | 18.57 | 17.72 | 37.10 | 27.35 | 12 | 7 |
| 514 | LEU | GLY | 3.88E-08 | 18.57 | 18.03 | 37.10 | 33.68 | 12 | 7 |
| 514 | LEU | ALA | 9.24E-07 | 18.57 | 18.09 | 37.10 | 34.25 | 12 | 7 |
| 529 | THR | HID | 6.68E-08 | 18.58 | 18.58 | 36.99 | -10.11 | 11 | 7 |
| 529 | THR | HIE | 6.68E-08 | 18.58 | 18.58 | 36.99 | -4.75 | 11 | 7 |
| 529 | THR | ASP | 2.37E-06 | 18.58 | 18.51 | 36.99 | 34.7 | 11 | 7 |
| 531 | TRP | PRO | 5.44E-07 | 18.51 | 16.76 | 37.07 | 0 | 1 | 7 |
| 505 | LEU | ALA | 5.28E-07 | 18.59 | 18.58 | 36.85 | 35 | 5 | 8 |
| 529 | THR | HIP | 6.68E-08 | 18.58 | 18.58 | 36.99 | -6.63 | 11 | 8 |

**Table S5:** All Resistor resistance mutation predictions for BRAF with vemurafenib. “Pos” is the position of the residue. “WT AA” is the wildtype identity of the amino acid. “Mut AA” is the resistance mutation. “Sig Prob” is the mutational signature probability for the mutation from “WT AA” to “Mut AA” in melanoma. “ATP WT” and “ATP Mut” are the *K*^*^ scores of the endogenous ligand with the wildtype and mutant residues, respectively. “Drug WT” and “Drug Mut” are the *K*^*^ scores of vemurafenib with the wildtype and mutant residues, respectively. “Count” is number of resistance mutations at the position. “Rank” is the Pareto rank of the mutation. Note: *K*^*^ scores are in log_10_ units where possible and 0 where there is predicted to be no binding.

| Pos | WT AA | Mut AA | Sig Prob | ATP WT | ATP Mut | Drug WT | Drug Mut | Count | Rank |
| --- | --- | --- | --- | --- | --- | --- | --- | --- | --- |
| 471 | VAL | LEU | 0.000443 | 18.65 | 19.67 | 33.41 | 29.39 | 4 | 1 |
| 481 | ALA | THR | 0.0177 | 18.58 | 18.99 | 33.27 | 30.69 | 9 | 1 |
| 529 | THR | ILE | 0.0202 | 18.58 | 18.57 | 33.45 | 29.33 | 10 | 1 |
| 535 | SER | PRO | 0.0013 | 18.72 | 18.65 | 33.65 | 0 | 1 | 1 |
| 593 | GLY | PHE | 1.99E-06 | 18.66 | 20.07 | 33.47 | 0 | 16 | 1 |
| 593 | GLY | TYR | 3.58E-05 | 18.66 | 19.86 | 33.47 | 0 | 16 | 1 |
| 593 | GLY | ARG | 0.00078 | 18.66 | 16.17 | 33.47 | -231.93 | 16 | 1 |
| 593 | GLY | ASN | 0.00134 | 18.66 | 19.16 | 33.47 | -103.95 | 16 | 1 |
| 593 | GLY | ASP | 0.0163 | 18.66 | 18.89 | 33.47 | -26.78 | 16 | 1 |
| 593 | GLY | CYS | 0.00166 | 18.66 | 19.06 | 33.47 | 19.89 | 16 | 1 |
| 593 | GLY | VAL | 0.000935 | 18.66 | 19.18 | 33.47 | 21.09 | 16 | 1 |
| 593 | GLY | ILE | 4.55E-05 | 18.66 | 19.8 | 33.47 | 23.08 | 16 | 1 |
| 593 | GLY | SER | 0.0609 | 18.66 | 18.83 | 33.47 | 31.33 | 16 | 1 |
| 463 | ILE | TYR | 2.55E-05 | 18.60 | 15.94 | 33.42 | -124.43 | 4 | 2 |
| 481 | ALA | GLU | 0.00121 | 18.58 | 17.92 | 33.27 | 11.12 | 9 | 2 |
| 481 | ALA | VAL | 0.00146 | 18.58 | 17.77 | 33.27 | 28.52 | 9 | 2 |
| 505 | LEU | PHE | 0.00544 | 18.59 | 18.58 | 33.45 | 27.01 | 3 | 2 |
| 505 | LEU | ARG | 0.000861 | 18.59 | 18.58 | 33.45 | 27.41 | 3 | 2 |
| 508 | THR | ARG | 0.000295 | 18.59 | 18.59 | 33.45 | 12.91 | 2 | 2 |
| 508 | THR | LYS | 0.000916 | 18.59 | 18.59 | 33.45 | 21.18 | 2 | 2 |
| 514 | LEU | ILE | 0.00165 | 18.57 | 18.4 | 33.43 | 30.08 | 11 | 2 |
| 532 | CYS | ARG | 0.000812 | 18.49 | 13.04 | 33.28 | -10.66 | 9 | 2 |
| 593 | GLY | HIE | 1.66E-05 | 18.66 | 19.58 | 33.47 | 0 | 16 | 2 |
| 593 | GLY | GLU | 0.000276 | 18.66 | 18.73 | 33.47 | -12.82 | 16 | 2 |
| 593 | GLY | THR | 2.67E-05 | 18.66 | 19.06 | 33.47 | 20.72 | 16 | 2 |
| 481 | ALA | LEU | 7.06E-05 | 18.58 | 18.68 | 33.27 | 18.19 | 9 | 3 |
| 481 | ALA | LYS | 0.000274 | 18.58 | 17.32 | 33.27 | 18.46 | 9 | 3 |
| 514 | LEU | ARG | 5.22E-05 | 18.57 | 17.18 | 33.43 | 22.28 | 11 | 3 |
| 514 | LEU | GLN | 8.77E-05 | 18.57 | 17.02 | 33.43 | 27.9 | 11 | 3 |
| 514 | LEU | SER | 0.000227 | 18.57 | 18.1 | 33.43 | 30.81 | 11 | 3 |
| 529 | THR | MET | 1.74E-05 | 18.58 | 18.65 | 33.45 | -8.52 | 10 | 3 |
| 529 | THR | PHE | 3.77E-05 | 18.58 | 15.91 | 33.45 | 5.09 | 10 | 3 |
| 593 | GLY | HIP | 1.66E-05 | 18.66 | 19.56 | 33.47 | 0 | 16 | 3 |
| 481 | ALA | ILE | 5.14E-05 | 18.58 | 14.2 | 33.27 | 24.57 | 9 | 4 |
| 593 | GLY | HID | 1.66E-05 | 18.66 | 19.55 | 33.47 | 0 | 16 | 4 |
| 471 | VAL | MET | 8.44E-06 | 18.65 | 18.96 | 33.41 | 31.15 | 4 | 5 |
| 481 | ALA | ARG | 1.53E-05 | 18.58 | 17.02 | 33.27 | 19.55 | 9 | 5 |
| 481 | ALA | ASP | 4.39E-06 | 18.58 | 19.21 | 33.27 | 24.15 | 9 | 5 |
| 514 | LEU | PHE | 1.53E-05 | 18.57 | 18.19 | 33.43 | 0 | 11 | 5 |
| 593 | GLY | TRP | 4.78E-06 | 18.66 | 18.83 | 33.47 | 0 | 16 | 5 |
| 593 | GLY | LEU | 3.87E-07 | 18.66 | 19.35 | 33.47 | -237.44 | 16 | 5 |
| 463 | ILE | HIE | 5.14E-06 | 18.60 | 18.06 | 33.42 | 30.11 | 4 | 6 |
| 463 | ILE | HID | 5.14E-06 | 18.60 | 18.08 | 33.42 | 30.75 | 4 | 6 |
| 466 | GLY | PRO | 1.77E-07 | 18.80 | 18.27 | 33.48 | 0 | 1 | 6 |
| 471 | VAL | PRO | 5.74E-07 | 18.65 | 17.88 | 33.41 | 0 | 4 | 6 |
| 471 | VAL | GLU | 9.01E-07 | 18.65 | 18.82 | 33.41 | 31.34 | 4 | 6 |
| 481 | ALA | GLN | 1.8E-06 | 18.58 | 18.37 | 33.27 | -15.45 | 9 | 6 |
| 514 | LEU | HIP | 3.37E-07 | 18.57 | 17.68 | 33.43 | 25.66 | 11 | 6 |
| 514 | LEU | HID | 3.37E-07 | 18.57 | 17.72 | 33.43 | 25.78 | 11 | 6 |
| 514 | LEU | GLY | 3.88E-08 | 18.57 | 18.03 | 33.43 | 30.29 | 11 | 6 |
| 514 | LEU | ALA | 9.24E-07 | 18.57 | 18.09 | 33.43 | 30.82 | 11 | 6 |
| 516 | PHE | ARG | 4.47E-06 | 18.59 | 18.58 | 33.51 | 29.7 | 1 | 6 |
| 529 | THR | TYR | 1.12E-06 | 18.58 | 18.58 | 33.45 | -114.94 | 10 | 6 |
| 529 | THR | ARG | 4.98E-07 | 18.58 | 18.6 | 33.45 | -34.39 | 10 | 6 |
| 529 | THR | LYS | 1.74E-06 | 18.58 | 18.57 | 33.45 | -7.92 | 10 | 6 |
| 529 | THR | LEU | 3.1E-06 | 18.58 | 18.56 | 33.45 | 28.04 | 10 | 6 |
| 532 | CYS | HIP | 4.44E-06 | 18.49 | 14.51 | 33.28 | -0.94 | 9 | 6 |
| 532 | CYS | HIE | 4.44E-06 | 18.49 | 12.48 | 33.28 | -2.8 | 9 | 6 |
| 532 | CYS | ILE | 1.16E-06 | 18.49 | 18.82 | 33.28 | 27.39 | 9 | 6 |
| 532 | CYS | VAL | 2.11E-06 | 18.49 | 18.7 | 33.28 | 29.58 | 9 | 6 |
| 463 | ILE | HIP | 5.14E-06 | 18.60 | 17.51 | 33.42 | 30.23 | 4 | 7 |
| 505 | LEU | MET | 3.58E-07 | 18.59 | 18.6 | 33.45 | 25.88 | 3 | 7 |
| 514 | LEU | HIE | 3.37E-07 | 18.57 | 17.72 | 33.43 | 25.91 | 11 | 7 |
| 529 | THR | HIE | 6.68E-08 | 18.58 | 18.58 | 33.45 | 10.11 | 10 | 7 |
| 531 | TRP | PRO | 5.44E-07 | 18.51 | 16.76 | 33.25 | 0 | 1 | 7 |
| 532 | CYS | HID | 4.44E-06 | 18.49 | 14.39 | 33.28 | -0.66 | 9 | 7 |
| 532 | CYS | THR | 2.67E-07 | 18.49 | 18.32 | 33.28 | 30.91 | 9 | 7 |
| 514 | LEU | GLU | 5.81E-08 | 18.57 | 17.36 | 33.43 | 28 | 11 | 8 |
| 529 | THR | HIP | 6.68E-08 | 18.58 | 18.58 | 33.45 | 10.86 | 10 | 8 |
| 529 | THR | HID | 6.68E-08 | 18.58 | 18.58 | 33.45 | 11.36 | 10 | 8 |

**Table S6:** All Resistor resistance mutation predictions for BRAF with encorafenib. “Pos” is the position of the residue. “WT AA” is the wildtype identity of the amino acid. “Mut AA” is the resistance mutation. “Sig Prob” is the mutational signature probability for the mutation from “WT AA” to “Mut AA” in melanoma. “ATP WT” and “ATP Mut” are the *K*^*^ scores of the endogenous ligand with the wildtype and mutant residues, respectively. “Drug WT” and “Drug Mut” are the *K*^*^ scores of encorafenib with the wildtype and mutant residues, respectively. “Count” is number of resistance mutations at the position. “Rank” is the Pareto rank of the mutation. Note: *K*^*^ scores are in log_10_ units where possible and 0 where there is predicted to be no binding.

| Pos | WT AA | Mut AA | Sig Prob | ATP WT | ATP Mut | Drug WT | Drug Mut | Count | Rank |
| --- | --- | --- | --- | --- | --- | --- | --- | --- | --- |
| 471 | VAL | LEU | 0.000443 | 18.65 | 19.67 | 38.16 | 28.68 | 10 | 1 |
| 481 | ALA | LEU | 7.06E-05 | 18.58 | 18.68 | 38.13 | -24.05 | 8 | 1 |
| 481 | ALA | GLU | 0.00121 | 18.58 | 17.92 | 38.13 | 10.6 | 8 | 1 |
| 529 | THR | ILE | 0.0202 | 18.58 | 18.57 | 38.12 | 31.23 | 12 | 1 |
| 532 | CYS | ARG | 0.000812 | 18.49 | 13.04 | 38.06 | -19.78 | 9 | 1 |
| 535 | SER | PRO | 0.0013 | 18.72 | 18.65 | 38.18 | 0 | 1 | 1 |
| 593 | GLY | TYR | 3.58E-05 | 18.66 | 19.86 | 38.09 | 0 | 17 | 1 |
| 593 | GLY | ARG | 0.00078 | 18.66 | 16.17 | 38.09 | -2.33 | 17 | 1 |
| 593 | GLY | ILE | 4.55E-05 | 18.66 | 19.8 | 38.09 | 2.23 | 17 | 1 |
| 593 | GLY | VAL | 0.000935 | 18.66 | 19.18 | 38.09 | 7.05 | 17 | 1 |
| 593 | GLY | ASN | 0.00134 | 18.66 | 19.16 | 38.09 | 28.19 | 17 | 1 |
| 593 | GLY | ASP | 0.0163 | 18.66 | 18.89 | 38.09 | 28.87 | 17 | 1 |
| 593 | GLY | PHE | 1.99E-06 | 18.66 | 20.07 | 38.09 | 30.05 | 17 | 1 |
| 593 | GLY | CYS | 0.00166 | 18.66 | 19.06 | 38.09 | 34.56 | 17 | 1 |
| 593 | GLY | SER | 0.0609 | 18.66 | 18.83 | 38.09 | 35.04 | 17 | 1 |
| 471 | VAL | PHE | 0.000785 | 18.65 | 16.37 | 38.16 | 26 | 10 | 2 |
| 481 | ALA | LYS | 0.000274 | 18.58 | 17.32 | 38.13 | 7.23 | 8 | 2 |
| 514 | LEU | ARG | 5.22E-05 | 18.57 | 17.18 | 38.14 | 20.73 | 10 | 2 |
| 529 | THR | MET | 1.74E-05 | 18.58 | 18.65 | 38.12 | -18.87 | 12 | 2 |
| 529 | THR | PHE | 3.77E-05 | 18.58 | 15.91 | 38.12 | 0.72 | 12 | 2 |
| 529 | THR | ASN | 0.000996 | 18.58 | 18.55 | 38.12 | 33.77 | 12 | 2 |
| 536 | SER | ASN | 0.0137 | 18.63 | 18.53 | 38.02 | 34.35 | 3 | 2 |
| 583 | PHE | TYR | 0.00408 | 18.60 | 18.68 | 38.10 | 29.81 | 9 | 2 |
| 593 | GLY | HIE | 1.66E-05 | 18.66 | 19.58 | 38.09 | 0 | 17 | 2 |
| 593 | GLY | GLU | 0.000276 | 18.66 | 18.73 | 38.09 | 22.97 | 17 | 2 |
| 593 | GLY | THR | 2.67E-05 | 18.66 | 19.06 | 38.09 | 24.19 | 17 | 2 |
| 593 | GLY | ALA | 0.000254 | 18.66 | 18.8 | 38.09 | 35.61 | 17 | 2 |
| 463 | ILE | TYR | 2.55E-05 | 18.60 | 15.94 | 38.08 | 9.07 | 3 | 3 |
| 481 | ALA | ILE | 5.14E-05 | 18.58 | 14.2 | 38.13 | 30.63 | 8 | 3 |
| 481 | ALA | VAL | 0.00146 | 18.58 | 17.77 | 38.13 | 34.23 | 8 | 3 |
| 505 | LEU | HIP | 0.00146 | 18.59 | 18.59 | 38.08 | 35.98 | 2 | 3 |
| 514 | LEU | GLN | 8.77E-05 | 18.57 | 17.02 | 38.14 | 31.64 | 10 | 3 |
| 536 | SER | ASP | 5.03E-05 | 18.63 | 18.21 | 38.02 | 33.54 | 3 | 3 |
| 583 | PHE | VAL | 0.000214 | 18.60 | 17.05 | 38.10 | 33.2 | 9 | 3 |
| 583 | PHE | ILE | 0.00316 | 18.60 | 17.45 | 38.10 | 34.49 | 9 | 3 |
| 583 | PHE | SER | 0.00262 | 18.60 | 16.86 | 38.10 | 34.31 | 9 | 3 |
| 593 | GLY | HIP | 1.66E-05 | 18.66 | 19.56 | 38.09 | 0 | 17 | 3 |
| 505 | LEU | HID | 0.00146 | 18.59 | 18.59 | 38.08 | 36.08 | 2 | 4 |
| 593 | GLY | HID | 1.66E-05 | 18.66 | 19.55 | 38.09 | 0 | 17 | 4 |
| 471 | VAL | PRO | 5.74E-07 | 18.65 | 17.88 | 38.16 | 0 | 10 | 5 |
| 471 | VAL | ARG | 3.03E-07 | 18.65 | 18.99 | 38.16 | 18.89 | 10 | 5 |
| 471 | VAL | MET | 8.44E-06 | 18.65 | 18.96 | 38.16 | 21.35 | 10 | 5 |
| 481 | ALA | GLN | 1.8E-06 | 18.58 | 18.37 | 38.13 | -7.07 | 8 | 5 |
| 481 | ALA | ARG | 1.53E-05 | 18.58 | 17.02 | 38.13 | 1.87 | 8 | 5 |
| 481 | ALA | ASP | 4.39E-06 | 18.58 | 19.21 | 38.13 | 30.87 | 8 | 5 |
| 514 | LEU | PHE | 1.53E-05 | 18.57 | 18.19 | 38.14 | -25.54 | 10 | 5 |
| 529 | THR | ARG | 4.98E-07 | 18.58 | 18.6 | 38.12 | -6.65 | 12 | 5 |
| 532 | CYS | HIP | 4.44E-06 | 18.49 | 14.51 | 38.06 | -40.5 | 9 | 5 |
| 532 | CYS | HIE | 4.44E-06 | 18.49 | 12.48 | 38.06 | -40.74 | 9 | 5 |
| 533 | GLU | PRO | 3.22E-07 | 18.58 | 18.63 | 38.12 | 0 | 1 | 5 |
| 593 | GLY | LEU | 3.87E-07 | 18.66 | 19.35 | 38.09 | 19.61 | 17 | 5 |
| 593 | GLY | TRP | 4.78E-06 | 18.66 | 18.83 | 38.09 | 19.1 | 17 | 5 |
| 463 | ILE | HIE | 5.14E-06 | 18.60 | 18.06 | 38.08 | 35.28 | 3 | 6 |
| 471 | VAL | GLU | 9.01E-07 | 18.65 | 18.82 | 38.16 | 35.77 | 10 | 6 |
| 529 | THR | TYR | 1.12E-06 | 18.58 | 18.58 | 38.12 | 19.14 | 12 | 6 |
| 529 | THR | LYS | 1.74E-06 | 18.58 | 18.57 | 38.12 | 20.27 | 12 | 6 |
| 529 | THR | HIE | 6.68E-08 | 18.58 | 18.58 | 38.12 | 21.51 | 12 | 6 |
| 529 | THR | LEU | 3.1E-06 | 18.58 | 18.56 | 38.12 | 22.42 | 12 | 6 |
| 531 | TRP | PRO | 5.44E-07 | 18.51 | 16.76 | 38.04 | 0 | 1 | 6 |
| 532 | CYS | HID | 4.44E-06 | 18.49 | 14.39 | 38.06 | -40.35 | 9 | 6 |
| 532 | CYS | ILE | 1.16E-06 | 18.49 | 18.82 | 38.06 | 30.25 | 9 | 6 |
| 532 | CYS | VAL | 2.11E-06 | 18.49 | 18.7 | 38.06 | 30.4 | 9 | 6 |
| 583 | PHE | ARG | 5.91E-06 | 18.60 | 17.17 | 38.10 | 32.32 | 9 | 6 |
| 583 | PHE | THR | 1.13E-05 | 18.60 | 16.91 | 38.10 | 32.78 | 9 | 6 |
| 583 | PHE | MET | 5.94E-06 | 18.60 | 18.39 | 38.10 | 35.48 | 9 | 6 |
| 463 | ILE | HID | 5.14E-06 | 18.60 | 18.08 | 38.08 | 35.51 | 3 | 7 |
| 471 | VAL | HIP | 6.58E-07 | 18.65 | 17.42 | 38.16 | 30.52 | 10 | 7 |
| 471 | VAL | HID | 6.58E-07 | 18.65 | 17.61 | 38.16 | 31.08 | 10 | 7 |
| 471 | VAL | TYR | 1.33E-06 | 18.65 | 16.37 | 38.16 | 31.46 | 10 | 7 |
| 514 | LEU | HIP | 3.37E-07 | 18.57 | 17.68 | 38.14 | 31.73 | 10 | 7 |
| 514 | LEU | HID | 3.37E-07 | 18.57 | 17.72 | 38.14 | 31.87 | 10 | 7 |
| 514 | LEU | LYS | 1.68E-06 | 18.57 | 17.37 | 38.14 | 33.97 | 10 | 7 |
| 514 | LEU | MET | 2.51E-06 | 18.57 | 18.42 | 38.14 | 35.5 | 10 | 7 |
| 529 | THR | HID | 6.68E-08 | 18.58 | 18.58 | 38.12 | 20.75 | 12 | 7 |
| 529 | THR | ASP | 2.37E-06 | 18.58 | 18.51 | 38.12 | 34.9 | 12 | 7 |
| 536 | SER | LEU | 5.47E-07 | 18.63 | 17.44 | 38.02 | 30.59 | 3 | 7 |
| 583 | PHE | TRP | 9.45E-07 | 18.60 | 13.08 | 38.10 | 24.17 | 9 | 7 |
| 583 | PHE | GLY | 1.52E-06 | 18.60 | 16.63 | 38.10 | 33.81 | 9 | 7 |
| 471 | VAL | HIE | 6.58E-07 | 18.65 | 17.28 | 38.16 | 30.93 | 10 | 8 |
| 514 | LEU | HIE | 3.37E-07 | 18.57 | 17.72 | 38.14 | 32.04 | 10 | 8 |
| 529 | THR | HIP | 6.68E-08 | 18.58 | 18.58 | 38.12 | 20.97 | 12 | 8 |
| 532 | CYS | THR | 2.67E-07 | 18.49 | 18.32 | 38.06 | 35.12 | 9 | 8 |
| 514 | LEU | GLU | 5.81E-08 | 18.57 | 17.36 | 38.14 | 32.45 | 10 | 9 |
| 514 | LEU | GLY | 3.88E-08 | 18.57 | 18.03 | 38.14 | 35.47 | 10 | 9 |

**Table S7:** All Resistor resistance mutation predictions for BRAF with PLX8394. “Pos” is the position of the residue. “WT AA” is the wildtype identity of the amino acid. “Mut AA” is the resistance mutation. “Sig Prob” is the mutational signature probability for the mutation from “WT AA” to “Mut AA” in melanoma. “ATP WT” and “ATP Mut” are the *K*^*^ scores of the endogenous ligand with the wildtype and mutant residues, respectively. “Drug WT” and “Drug Mut” are the *K*^*^ scores of PLX8394 with the wildtype and mutant residues, respectively. “Count” is number of resistance mutations at the position. “Rank” is the Pareto rank of the mutation. Note: *K*^*^ scores are in log_10_ units where possible and 0 where there is predicted to be no binding.

| Pos | WT AA | Mut AA | Sig Prob | ATP WT | ATP Mut | Drug WT | Drug Mut | Count | Rank |
| --- | --- | --- | --- | --- | --- | --- | --- | --- | --- |
| 471 | VAL | LEU | 0.000443 | 18.645 | 19.67 | 31.624 | 29.38 | 3 | 1 |
| 513 | ILE | PHE | 0.00146 | 18.569 | 18.57 | 31.663 | 0 | 1 | 1 |
| 529 | THR | ILE | 0.0202 | 18.583 | 18.57 | 32.196 | 12.61 | 14 | 1 |
| 535 | SER | PRO | 0.0013 | 18.717 | 18.65 | 31.813 | 0 | 5 | 1 |
| 535 | SER | LEU | 0.00156 | 18.717 | 19.09 | 31.813 | 24.04 | 5 | 1 |
| 593 | GLY | PHE | 1.99E-06 | 18.659 | 20.07 | 31.991 | 0 | 16 | 1 |
| 593 | GLY | TYR | 3.58E-05 | 18.659 | 19.86 | 31.991 | 0 | 16 | 1 |
| 593 | GLY | ARG | 0.00078 | 18.659 | 16.17 | 31.991 | -248.98 | 16 | 1 |
| 593 | GLY | GLU | 0.000276 | 18.659 | 18.73 | 31.991 | -61.35 | 16 | 1 |
| 593 | GLY | ASN | 0.00134 | 18.659 | 19.16 | 31.991 | -56.56 | 16 | 1 |
| 593 | GLY | ASP | 0.0163 | 18.659 | 18.89 | 31.991 | -29.16 | 16 | 1 |
| 593 | GLY | VAL | 0.000935 | 18.659 | 19.18 | 31.991 | 6.66 | 16 | 1 |
| 593 | GLY | CYS | 0.00166 | 18.659 | 19.06 | 31.991 | 12.77 | 16 | 1 |
| 593 | GLY | ILE | 4.55E-05 | 18.659 | 19.8 | 31.991 | 22.76 | 16 | 1 |
| 593 | GLY | SER | 0.0609 | 18.659 | 18.83 | 31.991 | 28.13 | 16 | 1 |
| 463 | ILE | TYR | 2.55E-05 | 18.596 | 15.94 | 31.621 | -260.1 | 5 | 2 |
| 481 | ALA | LEU | 7.06E-05 | 18.575 | 18.68 | 32.195 | 2.47 | 8 | 2 |
| 481 | ALA | LYS | 0.000274 | 18.575 | 17.32 | 32.195 | 3.05 | 8 | 2 |
| 481 | ALA | GLU | 0.00121 | 18.575 | 17.92 | 32.195 | 12.04 | 8 | 2 |
| 505 | LEU | ARG | 0.000861 | 18.589 | 18.58 | 31.429 | 24.96 | 4 | 2 |
| 508 | THR | ARG | 0.000295 | 18.594 | 18.59 | 31.461 | -115.96 | 2 | 2 |
| 508 | THR | LYS | 0.000916 | 18.594 | 18.59 | 31.461 | 12.12 | 2 | 2 |
| 514 | LEU | ILE | 0.00165 | 18.57 | 18.4 | 31.667 | 21.52 | 10 | 2 |
| 514 | LEU | ARG | 5.22E-05 | 18.57 | 17.18 | 31.667 | 21.09 | 10 | 2 |
| 529 | THR | PHE | 3.77E-05 | 18.583 | 15.91 | 32.196 | -106.19 | 14 | 2 |
| 529 | THR | MET | 1.74E-05 | 18.583 | 18.65 | 32.196 | -28.68 | 14 | 2 |
| 529 | THR | VAL | 4.99E-05 | 18.583 | 18.72 | 32.196 | 28.05 | 14 | 2 |
| 529 | THR | ASN | 0.000996 | 18.583 | 18.55 | 32.196 | 27.98 | 14 | 2 |
| 532 | CYS | ARG | 0.000812 | 18.49 | 13.04 | 31.578 | -0.71 | 9 | 2 |
| 535 | SER | ILE | 0.000391 | 18.717 | 19.02 | 31.813 | 28.17 | 5 | 2 |
| 535 | SER | TYR | 0.00262 | 18.717 | 18.68 | 31.813 | 29.19 | 5 | 2 |
| 593 | GLY | HIE | 1.66E-05 | 18.659 | 19.58 | 31.991 | 0 | 16 | 2 |
| 593 | GLY | THR | 2.67E-05 | 18.659 | 19.06 | 31.991 | 6.63 | 16 | 2 |
| 471 | VAL | PHE | 0.000785 | 18.645 | 16.37 | 31.624 | 27.32 | 3 | 3 |
| 481 | ALA | ILE | 5.14E-05 | 18.575 | 14.2 | 32.195 | 16.28 | 8 | 3 |
| 481 | ALA | VAL | 0.00146 | 18.575 | 17.77 | 32.195 | 27.92 | 8 | 3 |
| 514 | LEU | VAL | 0.000522 | 18.57 | 18.3 | 31.667 | 27.8 | 10 | 3 |
| 514 | LEU | PRO | 0.00121 | 18.57 | 18.12 | 31.667 | 29.31 | 10 | 3 |
| 593 | GLY | HIP | 1.66E-05 | 18.659 | 19.56 | 31.991 | 0 | 16 | 3 |
| 593 | GLY | HID | 1.66E-05 | 18.659 | 19.55 | 31.991 | 0 | 16 | 4 |
| 471 | VAL | MET | 8.44E-06 | 18.645 | 18.96 | 31.624 | 30.12 | 3 | 5 |
| 481 | ALA | ARG | 1.53E-05 | 18.575 | 17.02 | 32.195 | -7 | 8 | 5 |
| 481 | ALA | ASP | 4.39E-06 | 18.575 | 19.21 | 32.195 | 26.65 | 8 | 5 |
| 505 | LEU | TYR | 8.68E-06 | 18.589 | 18.59 | 31.429 | 13.95 | 4 | 5 |
| 514 | LEU | PHE | 1.53E-05 | 18.57 | 18.19 | 31.667 | 0 | 10 | 5 |
| 535 | SER | ARG | 1.91E-06 | 18.717 | 18.84 | 31.813 | 26.28 | 5 | 5 |
| 593 | GLY | TRP | 4.78E-06 | 18.659 | 18.83 | 31.991 | 0 | 16 | 5 |
| 593 | GLY | LEU | 3.87E-07 | 18.659 | 19.35 | 31.991 | -172.14 | 16 | 5 |
| 463 | ILE | HIE | 5.14E-06 | 18.596 | 18.06 | 31.621 | 14.78 | 5 | 6 |
| 463 | ILE | HIP | 5.14E-06 | 18.596 | 17.51 | 31.621 | 14.5 | 5 | 6 |
| 463 | ILE | HID | 5.14E-06 | 18.596 | 18.08 | 31.621 | 23.91 | 5 | 6 |
| 481 | ALA | GLN | 1.8E-06 | 18.575 | 18.37 | 32.195 | -57.06 | 8 | 6 |
| 516 | PHE | THR | 3.77E-06 | 18.593 | 18.56 | 32.276 | 29.08 | 1 | 6 |
| 528 | VAL | ARG | 5.57E-07 | 18.569 | 18.61 | 32.223 | 21.75 | 1 | 6 |
| 529 | THR | TYR | 1.12E-06 | 18.583 | 18.58 | 32.196 | 0 | 14 | 6 |
| 529 | THR | ARG | 4.98E-07 | 18.583 | 18.6 | 32.196 | -73.94 | 14 | 6 |
| 529 | THR | LEU | 3.1E-06 | 18.583 | 18.56 | 32.196 | -40.25 | 14 | 6 |
| 529 | THR | LYS | 1.74E-06 | 18.583 | 18.57 | 32.196 | -16.49 | 14 | 6 |
| 532 | CYS | HIP | 4.44E-06 | 18.49 | 14.51 | 31.578 | 10.43 | 9 | 6 |
| 532 | CYS | HIE | 4.44E-06 | 18.49 | 12.48 | 31.578 | 8.45 | 9 | 6 |
| 532 | CYS | ILE | 1.16E-06 | 18.49 | 18.82 | 31.578 | 28.7 | 9 | 6 |
| 532 | CYS | VAL | 2.11E-06 | 18.49 | 18.7 | 31.578 | 29 | 9 | 6 |
| 505 | LEU | MET | 3.58E-07 | 18.589 | 18.6 | 31.429 | 28.4 | 4 | 7 |
| 505 | LEU | GLN | 1.84E-06 | 18.589 | 18.6 | 31.429 | 29.15 | 4 | 7 |
| 514 | LEU | HIP | 3.37E-07 | 18.57 | 17.68 | 31.667 | 24.27 | 10 | 7 |
| 514 | LEU | HIE | 3.37E-07 | 18.57 | 17.72 | 31.667 | 24.93 | 10 | 7 |
| 514 | LEU | HID | 3.37E-07 | 18.57 | 17.72 | 31.667 | 25.66 | 10 | 7 |
| 529 | THR | HIP | 6.68E-08 | 18.583 | 18.58 | 32.196 | -75.31 | 14 | 7 |
| 529 | THR | HID | 6.68E-08 | 18.583 | 18.58 | 32.196 | -74.57 | 14 | 7 |
| 529 | THR | HIE | 6.68E-08 | 18.583 | 18.58 | 32.196 | -73.8 | 14 | 7 |
| 529 | THR | ASP | 2.37E-06 | 18.583 | 18.51 | 32.196 | 27.31 | 14 | 7 |
| 529 | THR | CYS | 1.13E-07 | 18.583 | 18.53 | 32.196 | 29.46 | 14 | 7 |
| 531 | TRP | PRO | 5.44E-07 | 18.512 | 16.76 | 31.57 | 0 | 1 | 7 |
| 532 | CYS | HID | 4.44E-06 | 18.49 | 14.39 | 31.578 | 10.57 | 9 | 7 |
| 514 | LEU | GLU | 5.81E-08 | 18.57 | 17.36 | 31.667 | 26.07 | 10 | 8 |
| 514 | LEU | LYS | 1.68E-06 | 18.57 | 17.37 | 31.667 | 28.06 | 10 | 8 |

## Footnotes

1 In the future, *c*_0_ could be learned from running Resistor on a resistance mutation dataset for homologous systems and examining the *K*^*^ scores.

